# Factors Underlying Asymmetric Dynamics of Disaggregase and Microtubule Severing AAA+ Machines

**DOI:** 10.1101/2020.11.25.398420

**Authors:** Mangesh Damre, Ashan Dayananda, Rohith Anand Varikoti, George Stan, Ruxandra I. Dima

## Abstract

Disaggregation and microtubule-severing nanomachines from the AAA+ (ATPases associated with various cellular activities) superfamily assemble into ring–shaped hexamers that enable protein remodeling by coupling large–scale conformational changes with application of mechanical forces within a central pore by loops protruding within the pore. We probed these motions and intra-ring interactions that support them by performing extensive explicit solvent molecular dynamics simulations of single-ring severing proteins and the double-ring disaggregase ClpB. Simulations reveal that dynamic stability of hexamers of severing proteins and of the nucleotide binding domain 1 (NBD1) ring of ClpB, which belong to the same clade, involves a network of salt bridges that connect conserved motifs of central PL1 loops of the hexamer. Clustering analysis of ClpB highlights correlated motions of domains of neighboring protomers supporting strong inter-protomer collaboration. Severing proteins have weaker inter-protomer coupling and stronger intra-protomer stabilization through salt bridges formed between PL2 and PL3 loops. Distinct mechanisms are identified in the NBD2 ring of ClpB involving weaker inter–protomer coupling through salt bridges formed by non–canonical loops and stronger intra–protomer coupling. Pore width fluctuations associated with the PL1 constriction in the spiral states, in the presence of a substrate peptide, highlight stark differences between narrowing of channels of severing proteins and widening of the NBD1 ring of ClpB. This indicates divergent substrate processing mechanisms of remodeling and translocation by ClpB and substrate tail-end gripping and possible wedging on microtubule lattice by severing enzymes. Relaxation dynamics of the distance between the PL1 loops and the centers of mass of protomers reveals observation-time-dependent dynamics, leading to predicted relaxation times of tens of microseconds on millisecond experimental timescales. For ClpB the predicted relaxation time is in excellent agreement with the extracted time from smFRET experiments.

## 1. Introduction

AAA+ (ATPases Associated with diverse cellular Activities) nanomachines couple chemical energy and mechanical action to assist a broad range of essential cellular mechanisms including protein quality control, such as protein degradation and disaggregation, membrane fusion, DNA replication, microtubule disassembly or cargo transport along microtubules.^1–3^ Members of the AAA+ class involved in protein remodeling, such as caseinolytic proteases (Clp) and eukaryotic heat shock proteins Hsp100, and in microtubule severing, such as spastin, katanin and fidgetin, assemble into hexameric rings that encompass a narrow pore. During ATP-driven allosteric cycles, flexible central channel loops from each nucleotide binding domain (NBD) grip the substrate and mediate the application of mechanical force to promote unfolding or disassembly mechanisms.^4^ Class 1 proteins, such as ClpA, ClpB/Hsp104, and ClpC, have two NBDs per protomer that give rise to double–ring structures, whereas class 2 proteins, such as ClpX and microtubule-severing proteins, have a single NBD per protomer and form single–ring structures.^5,6^ From an evolutionary perspective, the NBD rings of class 1 proteins belong to distinct subgroups of the AAA+ superfamily: the NBD1 ring is part of clade 3 along with severing proteins and the NBD2 ring is part of clade 5 along with ClpX. The NBD consists of the highly conserved Walker–A (WA), Walker–B (WB), Arg–finger (RR) motifs and central pore loops.

Intriguingly, a large number of recent structural studies have revealed non–planar arrangements of protomers in the hexameric structures of these machines resulting in “spiral” or “ring” conformations.^7–31^ Protomers on each side of the oligomeric seam are found in distinct nucleotide states and display large conformational differences between the two configurations, therefore they are attributed active roles in promoting the mechanical action, whereas the other protomers are proposed to support hexamer stability and substrate gripping. Functionally, conformational asymmetry of Clp ATPases is proposed to underlie either sequential^16,30–32^ or probabilistic^28,33–39^ substrate gripping and translocation mechanisms, whereas its role in severing mechanisms is less clear. In addition, dynamic dissociation of the hexamer is proposed to act as a mechanism for releasing substrates targeted for degradation that are trapped into configurations that require excessively long processing times^40,41^ or for disengaging severing proteins from microtubules. ^39^ Currently, it is not well understood how these structural and functional aspects are enabled by hexamer dynamics and inter– protomer interactions. The variety of proposed mechanisms, formulated on the basis of the solved cryo–EM structures and biophysical and biochemical studies, calls for an understanding of the basis of the structural flexibility and of the dynamics of the various hexameric forms of disaggregases or severing enzymes. Moreover, in light of the proposed instability of the hexamers upon ATP hydrolysis and/or disengagement from the substrate peptide, it is important to probe the sources of structural stability in these oligomers.

Microtubule severing enzymes, such as katanin and spastin, influence various aspects of cellular dynamics such as meiosis, mitosis, ciliogenesis, and neuronal morphogenesis through their action on microtubules. ^42^ Moreover, mutations in these enzymes are associated with neurological disorders. Microtubules are the longest and most rigid elements of the cytoskeleton and are composed of polymeric assemblies of alpha– and beta–tubulin dimers linked by non–covalent longitudinal bonds into protofilaments that assemble laterally to form the microtubule lattice. The textbook function of the severing enzymes is to destabilize microtubules by binding along the lattice and inducing the fragmentation of the filament. ^39,43,44^ Recent experiments found that severing enzymes can also serve as amplifiers of microtubule arrays if the newly cut microtubules are stable and able to grow through tubulin exchange along the lattice shaft. ^39,45^ Severing enzymes oligomerize into hexamers in the presence of ATP and of negatively charged intrinsically disordered carboxy–terminal tails (CTTs) of the tubulin subunits that project from the microtubule surface, which constitute their primary binding sites on microtubules. Interestingly, studies revealed differences between katanin and spastin. Whereas both spastin and katanin can sever microtubules, only katanin can catalyze microtubule depolymerization in vitro and in vivo. ^46^ Furthermore, katanin can bind to microtubules or tubulin dimers specifically and tightly, and its severing activity is concentration dependent, ^47^ whereas spastin binds exclusively to microtubules, and the polyglutamylation of CTT side chains amplifies its severing activity. ^48^ Therefore, it is important to study both severing enzymes to understand their distinct actions on microtubules.

The ATPase domain in severing proteins comprises a large NBD domain and a smaller four–helix bundle domain (HBD). The NBD forms the central pore for substrate binding and the HBD forms a semi crescent structure covering the NBD region of the adjacent protomer. Protomers B through E make canonical convex–to–concave AAA interactions between successive protomers, i.e., both NBD and HBD from protomer *i* interact with the NBD from protomer i–1. The NBD in severing proteins has three central pore loops (PL1, PL2, and PL3). HBD has 2 functionally important regions: the sensor II motif and the C– Hlx.^39^ The central pore loops PL1 and PL2 from all 6 protomers in the spiral conformation are arranged in a double–spiral structure oriented from A to F around the poly–glutamate substrate used in the cryo–EM experiments. ^30^ Salt bridges formed by positions in the pore loops have been identified, which are important for the functions of the machine: ^29,30^ in the katanin spiral the intra–protomer salt bridge R267–E308, which connects pore loops PL1 and PL2, and in the spastin spiral the inter–protomer salt bridge R600–E633, which connects pore loops PL2 and PL3. Mutations in these positions led to substantial loss of ATPase activity and loss of severing function.

ClpB is distinct among members of the Clp ATPase family through its lack of association with a peptidase compartment and its ability to function as a disaggregase to provide protection against thermal stress. In collaboration with the DnaK/DnaJ/GrpE system, it rescues proteins from toxic aggregates^49–52^ and its engineered variant is able to promote protein degradation by threading substrates to an associated ClpP peptidase as do ClpX or ClpA. ^53^ In vivo, both a full–length and a N–terminal domain truncation variant are able to promote disaggregation ^54^ and remodeling activity can be elicited by a mixture of ATP and ATPqS to enable disaggregation or unfolding independent of cochaperones.^55^ Canonical pore loops of the two ClpB NBD domains, PL1 and PL3, actively participate in engaging the substrate and promoting mechanical action, however they are suggested to have non–overlapping functional roles. Whereas PL1 loops are involved in substrate recognition and are proposed to collaborate in stabilizing substrate engagement, PL3 loops operate non–cooperatively in DNA–dependent disaggregation.^56^ Each canonical pore loop contains a highly conserved Tyrosine residue which has the major contribution in substrate gripping and translocation. The PL1 loop within NBD1 contains the conserved KYR motif, homologous to those present in corresponding loops of severing proteins, which allows formation of salt bridges with the neighboring protomer. Cryo–EM experiments highlighted cross–protomer interactions involving residues K250 and R252 of one protomer and residues E254 and E256 of neighboring protomers.^25^ The resulting salt bridge network is proposed to support the stability to the hexameric structure. The PL3 loop within the NBD2 domain contains the conserved motif GYVG present in ClpX and ClpA loops which clamps around the substrate backbone and provides strong interaction and grip during the substrate translocation. Non–canonical pore loops, PL2 in NBD1 and PL4 in NBD2, which extend into the central pore of the hexameric structure form an additional spiral to provide support for interaction with the substrate.

Computational studies revealed strong effects of the pore conformation and dynamics on substrate remodeling. Simulations using model pores indicate dramatically different unfolding requirements of SPs near the pore entrance compared with bulk mechanical unfolding.^57–60^ In addition, simulations using molecular–level representation and details of allosteric cycles of the ATPase reveal that conformational plasticity of the pore surface provides a selectivity filter for SP orientations that favor application of mechanical force along weaker mechanical directions.^61–63^ In accord with experimental studies, simulations highlight kinetic effects that result from transient binding and release of SPs through kinetic competition between SP refolding and translocation.^4,64,65^

In this paper, we compare and contrast the dynamics and inter–protomer interactions of microtubule–severing proteins, spastin and katanin, and the ClpB disaggregase. To this end, we perform molecular dynamics (MD) simulations of fully solvated systems of each of these nanomachines in configurations that correspond to the nucleotide– and substrate–bound states present in cryoEM experiments. In addition, for each system, we probe the effect of removal of nucleotide from the active protomers and/or of the substrate on hexamer dynamics and inter–protomer interactions. The combined total duration for our extensive simulations is 6 μs. We find that severing proteins and the NBD1 ring of ClpB are dynamically stabilized by networks of cross–protomer complex salt bridges that involve residues in the conserved motif of PL1 loops. Clustering analysis of domain motions highlights inter–protomer coupling that supports strong collaboration within the hexameric ring. By contrast, the NBD2 ring of ClpB is characterized by weaker interactions between PL3 loops and predominantly intra– protomer coupling of domain motions, which supports non–cooperative models of interaction between domains in this ring and the substrate. Similarly, we found that a large network of intra–protomer salt bridges connects PL2 and PL3 loops in the ring states of the severing proteins, underlying the reduced interactions between these loops and the substrate. Our analysis of the relaxation dynamics of the distance between the PL1 loops and the center of mass of each protomer yields results indicative of the coupling between collective and local motions and observation–time–dependent dynamics. On the basis of these results, we predict relaxation times on the order of tens of microseconds on the millisecond timescales of single–molecule (smFRET) experiments.^66^ Comparison of the predicted relaxation times of ClpB with the corresponding time extracted from smFRET experiments that probed conformational dynamics of ClpB pore–loops ^67^ reveals excellent agreement between the two values.

## 2. Materials and Methods

### 2.1. Initial configuration

We used Protein Data Bank (PDB)^68^ structures, obtained using 3D cryo–EM, of katanin and spastin in two different conformations, spiral and ring, solved in the presence of minimal substrates and nucleotides (Table S1).^29–31^ Initial conformations of ClpB were obtained from the PDB structures 6OAX (ring) and 6OAY (spiral), which were solved using cryo–EM, in the presence of minimal substrate and nucleotides (Table S2). ^25^

We considered the following configurations of ClpB and severing enzymes: (i) substrate– free and nucleotide–free ATPase, termed APO; (ii) substrate–bound (polyE in 6UGD, 6UGE, 6P07, poly(EY) peptide in 6PEN, and polyA in 6OAX, 6OAY) and nucleotide–free ATPase, termed E14, E12, E15, or (EY)5 for severing proteins and ALA for ClpB; (iii) substrate–free ATPase and nucleotide–bound to all protomers according to the cryo–EM structure: with ATP in all 6 protomers from the spiral states and in 5 of the 6 protomers (exception is protomer A from 6UGE) in severing proteins, termed ATP and ADP in 5 of the 6 protomers (exception is protomer F from spastin ring 6PEN), termed ADP, and ATP (ATPqS) and ADP in 6OAX and 6OAY for ClpB, termed ATP; (iv) substrate–bound and nucleotide–bound ATPase, termed ATP+E14 (E12, E15, (EY)5) for severing proteins and ALA+ATP for ClpB. Details regarding the preparation of the configurations are provided in the Supplementary Material.

### 2.2. Molecular Dynamics Simulations

We performed MD simulations using the GROMACS ^69^ molecular modeling program version 2019 and the GROMOS96 54a7 force field.^70^ Five 50 ns production trajectories were obtained for each of the above configurations (Table S3 and S4). Details of the simulation procedures and parameters used are in the Supporting Information.

## 3. Data Analysis

### 3.1. RMSF and Free Energy Landscape

To determine the contribution of each amino acid to the motion of our proteins, we calculated the root mean square fluctuation (RMSF) of C_α_ atoms of each residue over MD trajectories of a given setup, as detailed in the Supplementary Information.

We used the free energy landscape (FEL) analysis to determine the extent of the conformational changes occurring in the central pore of the hexamer during our simulations as a result of the alteration in the binding partners. For severing proteins, we calculated the FEL in the PL1 and RMSD plane, and the FEL in the PL2 and RMSD plane. The FEL isgiven by:

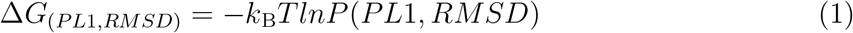

where *k*_B_ and *T* are the Boltzmann constant and absolute temperature, respectively, and *P(PL1, RMSD)* is the probability distribution of the molecular system along the PL1 and RMSD coordinates. PL1 here stands for the distance between diagonal PL1 loops (A–D, B–E, or C–F) that shows the largest degree of change in between the different setups for a given hexameric state (spiral or ring). For the FEL based on the PL2 distances, we used the distance between the same pairs of protomers used for the FEL based on the PL1 distance. For katanin, the PL1 distance is the distance between the C_α_ atoms of the invariant R267 in the PL1 loops from diagonally opposite protomers: C and F in the spiral state, and A and D in the ring state. For the spastin spiral, the PL1 distance is the distance between the C_α_ atoms of the invariant Y556 in the PL1 loops from the diagonally opposite C and F protomers. For the spastin ring, the PL1 distance is the distance between the C_α_ atoms of the invariant Y415 in the PL1 loops from the diagonally opposite C and F protomers. For katanin, the PL2 distance is the distance between the C_α_ atoms of the invariant H307 in the PL2 loops. For spastin spiral, the PL2 distance is the distance between the C_α_ atoms of the invariant H596 in the PL2 loops. Finally, for spastin ring, the PL2 distance is the distance between the C_α_ atoms of the invariant H455 in the PL2 loops. In the case of ClpB, we probed the combined probability distribution of the NBD1 (NBD2) pore width and backbone RMSD. As for severing proteins, the pore width is characterized by the diagonal distances of PL1 (PL3) pore loops between opposite i, i+3 protomers of the hexamer (A–D, B–E and C–F). For the FEL based on PL1 distances, we used the distance between the C_α_ atoms of the Y251 residues of B–E diagonal PL1 loops which showed the largest variation during the period of analysis. For the PL3 distances, we probed the B–E diagonal distance between C_α_ atoms of Y653 residues.

### 3.2. Dynamic Cross–Correlation Analysis

To discern correlated motions of regions in the hexameric structures during our MD simulations, we employed the correlation network analysis (CNA) implemented in the Bio3D package. ^71^ This approach uses the dynamic cross correlation maps (DCCMs) that quantify directional coupling of residue pairs according to

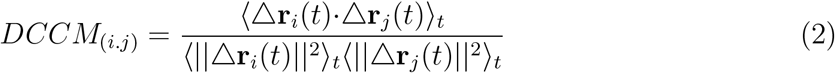

where **r**_*i*_*(t)* and **r**_*j*_*(t)* are the position vectors of C_α_ atoms of residues *i* and *j*, 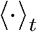, is the time ensemble average, and 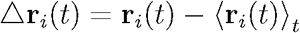, 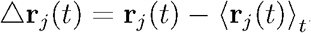. Motions obtained from DCCMs are highly interconnected at residue level and thus further analysis is needed to determine which regions of a hexamer work together in each state. To this end, DCCMs are used to build cross–correlation networks that highlight inter–residue connectivity. The resulting residue network is then converted into coarse–grained community network using the Girvan–Newman clustering method. ^71,72^

### 3.3. Salt bridge analysis

Salt bridges (SB) are a close range electrostatic interaction between a positively charged (Arg, Lys or His) and a negatively charged residue (Asp or Glu) as an ion pair.^73^ Salt bridges were calculated using the Visual Molecular Dynamics^74^ program with the criterion of a maximum distance of 4 A between at least one atom pair formed by a side chain carbonyl oxygen atom in the acidic residue and a side chain nitrogen atom in the basic residue. We monitored several salt bridges formed by amino acids located in the central pore region of the hexamer that persist through at least 1 ns during a simulation trajectory,

### 3.4. Relaxation Times

We determined the time dependence of structural fluctuations in our hexamers by calculating, for each protomer, the characteristic relaxation time of the distance between its PL1 loop and its center of mass. Namely, we extracted the autocorrelation function (ACF) of the distance between the central highly conserved PL1 residue (R267 in katanin, Y556 in spastin spiral, Y415 in spastin ring and R252 in ClpB) from each chain and the center of mass of the respective protomer using the Gromacs tool ‘analyze’. To make sure that we are only evaluating the internal fluctuations of the protein, prior to the calculation of the ACF we removed rigid–body translation and molecular rotations, using the ‘analyze’ subroutine. The time decay of the ACF is complex and non–exponential: for short time intervals (ps) we fitted it with a stretched exponential, using the Kohlrausch-Williams-Watts (mKWW) function. ^75^ To model the full decay of the ACF relaxation over tens of nanoseconds, we used the combination between a single–exponential and the stretched-exponential model proposed in the literature^75,76^ given by

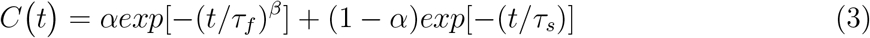

where *C(t)* is the relaxation function, *t* is the time, and *τ_f_* and *τ_s_* are the corresponding characteristic timescales from the fast and the slow processes, respectively and *β* is the dimensionless stretching exponent. For 0 ≤ β ≤ 1, the stretched exponential is the superposition of various timescale processes that contribute to the relaxation phenomena, each of which can be described by a single-exponential decay. The positive amplitude, 0 ≤ *α ≤* 1, accounts for the relative contributions of processes at short and long timescales.

The characteristic relaxation time of the autocorrelation function, τ*, is the mean relaxation time of the total decay,^75^ which based on (Eq. 3) is given by

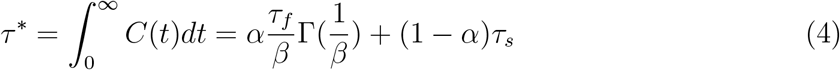

where we use the gamma function, Γ(1/β).^77^ We applied this fitting methodology to the ACF calculated from all simulation times (trajectory lengths): t = 5, 25, and 50 ns. For each trajectory length we used the function from (Eq. 3) only in the time region where *ACF* ≥ 0. Using Eq. 4, we calculated the respective characteristic relaxation time, *τ*^*^, for each protomer in each trajectory length and each simulation setup. For an individual protomer, we selected the characteristic *τ*^*^ value such that it corresponds to the time where the distribution of the χ^2^ values versus time for the fitting according to Eq. 3 has an elbow followed by a sharp increase in value.

## 4. Results and discussion

### 4.1. RMSF Analysis

Residue–wise averaged RMSF over the trajectories run for the various setups show differences in the backbone atomic fluctuations between the states of katanin and spastin (Fig. 1a) and S1). In the spiral states of both katanin and spastin (Fig. 1 and S1-b), we found that most of the protomers have the smallest fluctuations in the full spiral state (with ATP and substrate) and largest in the APO setup. In all protomers, with the exception of protomer A, the HBD region has the largest RMSF values, which can be assigned in part to its exposure to the solvent. The highest RMSF values for any region and any protomer are reached in the sensor II motif^39^ from the HBD of protomer F, irrespective of the state: 12 A in katanin and 8 A in spastin, compared to 5 Å for the next highest RMSF. We also found that the loop between PL2 and PL3 undergoes large fluctuations in many protomers, which we again assign to its high solvent accessibility. For example, the largest RMSF value in protomer A (5 Å) corresponds to this loop. In contrast, the RMSF values for the three functional pore loops are modest. Namely, PL1 and PL3 have very low RMSF values (up to 2.5 Å) in all protomers and in all cases. Interestingly, for PL2 loops we found cases in some protomers when the fluctuations go above 2.5 Å, but even these values are modest (up to 4 Å).

**Figure 1:**
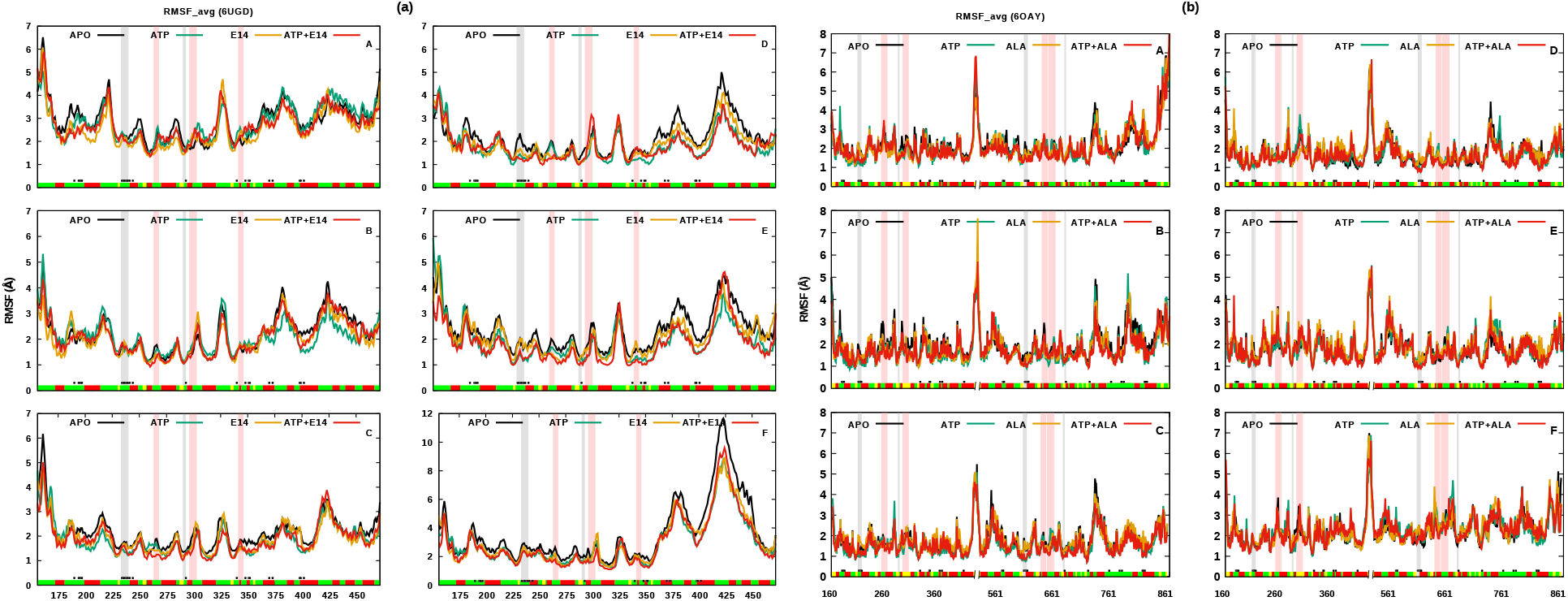
Average RMSF values for protomers in the spiral state of (a) katanin (6UGD) and (b) ClpB (6OAY), with secondary structures shown below each figure. For katanin we show the data in the APO state (black), ATP bound state (green), substrate bound state (orange) and substrate with ATP bound state (red). Different groups are annotated: black dots (·) are the ATP binding sites, and shaded regions show functionally important elements: WA and WB (in light gray), and PL1, PL2 and PL3 (in light red). For ClpB we show data in the APO state (blue), substrate-bound (ALA, orange), nucleotide–bound (ATP, green) and nucleotide– and substrate–bound (ALA–ATP, red).Different groups are annotated: black dots (·) are the ATP binding sites, and shaded regions show the functionally important elements: Pore loops (PL1–4) are in light red, Walker A (WA1–2) and B (WB1–2) regions are in light gray. The sequence gap between NBD1 and NBD2 domains is also indicated.

In the ring states of the severing proteins we found larger RMSF values (up to 5 Å) in the PL1 loops compared to the spiral case (Fig. S1-a and S1-c): PL1 has the highest fluctuations for the protomer (A in katanin and F in spastin) that lacks the nucleotide (ATP or, respectively, ADP) in the setups that include the substrate peptide. The highest value (5 Å) is reached in spastin. We thus conclude that for both katanin and spastin the stability of PL1 is dependent on the presence of the nucleotide (ATP or ADP). In contrast, PL2 and PL3 behave the same as in the spiral case. Similar to the spiral case, the largest RMSF values are in the sensor II motif of the HBD region for the states corresponding to either the presence of the minimal substrate or to the full ring state. The highest values are seen in the 2 end protomers: A and F. Interestingly, whereas in A the RMSF for the sensor II motif is higher than in the spiral case, in protomer F this is less than half of the value found in the corresponding spiral state (5 Å compared to 12 Å for katanin and 3.5 Å compared to 8 Å for spastin).

In summary, we found that large fluctuations in the sensor II motif of the HBD from the last protomer (F) are a signature of the severing proteins in their spiral state, with chain F being substantially more flexible in katanin than in spastin. This level of fluctuations in the sensor II motif recall the recent finding that, upon transition from an ADP–bound pseudo– hexameric state to a monomeric form the HBD in katanin exhibits refolding rearrangements in which sensor II elements are reorganized. ^39^ Moreover, the HBD region is usually the most flexible part in all the protomers in the spiral state. We also found that the 3 pore loops (PL1, PL2, and PL3) are rigid in most of the protomers in the presence of a minimal substrate and/or ATP molecules, which is likely an indication that the conformation of these loops needs to be maintained over time in order to engage the substrate. Because in the ring state PL1 is highly flexible in the protomer lacking the nucleotide, we propose that the binding of the nucleotide (ATP or ADP) in a protomer is required for the stability of PL1. As seen below in the FEL analysis, the close proximity of the PL1 loops in the various protomers is a signature of the spiral state in the presence of the substrate. On the basis of these findings, we conclude that the preservation of the conformation and the relative orientation of the pore loops from the majority of the protomers, coupled with their movement within close proximity from each other, are required for a severing machine to engage a substrate peptide. The increased flexibility of PL2 in half of the protomers in the spiral case when the substrate is present suggests that PL2 is designed to adapt to the presence of the substrate peptide.

RMSF analysis was also performed for backbone atoms in individual protomers of spiral and ring structures of ClpB in order to study their local conformational flexibility for all four configurations (APO, nucleotide–bound, substrate–bound, and nucleotide- and substrate– bound). As shown in Fig. 1b and S2, in both spiral and ring states, RMSF values of residues indicate a strong contrast between highly dynamic APO conformations and stabilized nucleotide– and substrate–bound ones. In the ring structure the largest RMSF values correspond to the seam protomer F and to the C–terminal region (residues 740–857) of protomer A, which have fluctuations larger than 2.5 Å. These observations are consistent with the weak connectivity of protomer F to the rest of the ClpB hexamer and with the ADP– bound nucleotide state of protomers A and F, which endows them with greater flexibility than the ATP–bound state of the other 4 protomers. Thus, ATP–binding to a subset of protomers results in dampening their conformational fluctuations, which helps to stabilize the hexameric structure and to provide support for the seam protomers that are actively promoting substrate translocation. In accord with these observations, we note that PL1 (residues 247–258) and PL2 (residues 284–295) regions of NBD1 have larger RMSF values in the ring state when the substrate is not present (APO and ATP), which indicates that these loops are actively engaged in substrate-gripping. We found that conformational fluctuations are lower in PL3 (residues 647–660) and PL4 (residues 636–646) regions of NBD2, except in protomers E and F, compared to PL1 and PL2. These differences can be attributed to the greater solvent accessibility of PL1 and PL2, which are located near the pore entrance, compared with PL3 and PL4 loops, which are located entirely within the ClpB channel. Greater flexibility of NBD2 loops of protomer F is in accord with their role in substrate translocation and with observations for the ring structure. The spiral structure involves larger fluctuations (> 2.5 Å) than the ring structure and RMSF values of residues in protomers A and F are more pronounced. Greater flexibility present in the spiral structure, particularly in the seam protomers A and F, is consistent with the initiation of the power cycle in this state. We also found that loop regions outside the seam have significantly lower RMSF values in the spiral state compared with the ring state, which indicates that strong gripping of the substrate by non–seam protomers is needed to assist substrate translocation. Notably, ClpB residues are found to have RMSF values of around 2.5 Å, which are generally significantly lower than those of katanin and spastin. Weaker ClpB flexibility can be attributed to its double–ring architecture that results in stabilizing hexameric interactions and in dampening of conformational fluctuations compared with single–ring katanin and spastin structures. Overall, we surmise that the double–ring structure is characterized by strong hexameric stability, especially associated with the non-seam protomers that support substrate gripping, and its larger conformational flexibility is associated with seam protomers that perform the translocation function.

### 4.2. Salt bridge analysis reveals dynamically stable cross–protomer networks of electrostatic interactions in both severing proteins and ClpB

Table 1 summarizes the number of salt bridges observed in both spiral and ring states of the severing proteins and Supplementary Table S5 indicates the specific salt bridge pairs observed with the highest frequency in our simulations. For katanin the number of the inter protomer salt bridges exceeds that of the intra protomer salt bridges in both the spiral and ring states (Table 1). This suggests that the overall organization and the stability of the pore loops in katanin is primarily due to the formation of specific electrostatic interactions resulting from the structural flexibility of the hexamer, rather than from localized motions inside a protomer. For spastin we found different patterns in the spiral and the ring states. In the spiral state of spastin, especially in the absence of ATP or of E15, the numbers of inter and intra protomers salt bridges are equal and are comparable to the number of inter protomer salt bridges in the spiral katanin structure. Thus, the stability of the central pore loops in the spastin spiral results from a combination of internal protomer motions and hexamer–wide conformational fluctuations. For the spastin ring the number of intra protomer salt bridges is higher than that of the inter protomer salt bridges. These are also the highest numbers of salt bridges found in our simulations, indicating that the pore loops in the spastin ring are stabilized primarily by salt bridges formed due to internal motions in the protomers.

**Table 1:** The average number of inter– and intra–protomer salt bridges that occur for at least 1 ns in MD trajectories of katanin, spastin, and ClpB proteins

| Spiral | State | Total |  | Ring | State | Total |  |
| --- | --- | --- | --- | --- | --- | --- | --- |
|  |  | Inter SB | Intra SB |  |  | Inter SB | Intra SB |
| **6UGD** | APO | 18±2 | 13±2 | **6UGE** | APO | 16±3 | 13±1 |
| | E14 | 17±1 | 10±2 | | E12 | 14±1 | 10±2 |
| | ATP | 17±2 | 12±2 | | ATP | 16±4 | 13±1 |
| | E14+ATP | 15±3 | 12±4 | | E12+ATP | 13±1 | 12±2 |
| **6P07** | APO | 14±3 | 14±1 | **6PEN** | APO | 23±2 | 27±4 |
| | E15 | 14±2 | 15±2 | | (EY)5 | 19±2 | 30±3 |
| | ATP | 13±2 | 15±2 | | ADP | 24±2 | 26±2 |
| | E15+ATP | 13±2 | 16±3 | | (EY)5+ADP | 21±3 | 24±4 |
|  |  | Total Inter SB |  | Total Intra SB |  |  |  |
| Spiral | State | NBD1 | NBD2 | NBD1 | NBD2 |  |  |
| **6OAY** | APO | 13±2 | 5±1 | 7±2 | 3±2 | | |
| | ALA | 11±2 | 4±1 | 8±1 | 5±1 | | |
| | ATP | 11±2 | 5±2 | 7±2 | 5±1 | | |
| | ALA+ATP | 12±1 | 5±0 | 6±1 | 5±1 | | |
| Ring | State | NBD1 | NBD2 | NBD1 | NBD2 |  |  |
| **6OAX** | APO | 12±1 | 5±1 | 7±3 | 6±1 | | |
| | ALA | 13±1 | 3±1 | 5±1 | 5±0 | | |
| | ATP | 12±2 | 4±1 | 6±1 | 4±1 | | |
| | ALA+ATP | 9±1 | 4±0 | 6±3 | 5±2 | | |

Next we concentrated on the salt bridges that are particularly stable in our simulations listed in Table S5. In contrast to the behavior of salt bridges in monomeric proteins, which are customarily found between residues separated by less than 10 positions in the sequence,,^73^ the most stable intra protomer salt bridges found in our simulations are formed between very distant positions (> 20). Importantly, in all the hexameric states of the severing proteins we found networks of positions from the 3 pore loops involved in the formation of complex salt bridges (between at least three charged residues) with each other. For the katanin spiral, the first position is K265 from PL1, which participates in 2 inter–protomer salt bridges in all states: with D171 from the fishhook and with D269 from PL1. These salt bridges are usually formed between 5 pairs of neighboring protomers. The mutation K265A led to a reduction in the basal and the MT stimulated ATPase activity, while completely inactivating severing. ^30^

Second, position R301 from PL2 participates in 2 salt bridges in all states: 1 inter–protomer with E293 from the Walker B motif, found usually between 5 pairs of neighboring protomers, and 1 intra–protomer with D346 from PL3, which is found in all 6 protomers. The mutation R301A reduced the ATPase activity and abolished microtubule severing.^30^ Third, position D346 from PL3 participates in 2 intra–protomer salt bridges: with R301 from PL2 in all protomers and with K314 usually in 4 protomers (the only exception is in the full ATP+E14 state, where it is found in all 6 protomers). Finally, position D269 from PL1 forms 2 salt bridges in the E14 and the ATP+E14 states: 1 inter–protomer with K265 from PL1 in 5 pairs of neighboring protomers and 1 intra–protomer with K272 in all protomers. Interestingly, the inter–protomer salt bridge identified based on the cryo–EM structure of the katanin spiral, ^30^ R267-E308, which has been proposed as important for the structuring of PL1 and PL2 loops, is not part of a network and neither residue forms complex salt bridges. We found it in the majority of the protomers in all states, with the exception of the one with just the substrate (E14) bound. For the katanin ring we found that the same set of 4 special positions are able to form a network of salt bridges as in the spiral state.

For the spastin spiral we also found 4 positions that participate in multiple salt bridges. Position R591 from PL2, which has been proposed as the center of an allosteric network that can couple the substrate binding loops to ATP hydrolysis and oligomerization, ^29^ participates in 3 salt bridges in the APO and ATP+E15 states: 2 inter–protomer (R591-D585 and R591-E633) present in 5 pairs of protomers, and 1 intra–protomer (R591-D635) present in all the protomers. D585 is in the Walker B motif so the first inter–protomer salt bridge is likely involved in the ATPase action of spastin, whereas E633 and D635 are in PL3. Importantly, R591 from *D. melanogaster* is the equivalent of R301 from *C. elegans* katanin (see Fig. 2E in^31^). Because R301 in katanin also forms multiple salt bridges, it is likely the center of an allosteric network that couples the substrate binding loops to ATP hydrolysis and oligomerization. The second residue, which participates in multiple salt bridges in the APO and the ATP+E15 states from 5 protomer pairs, is K555 from PL1: it forms 2 inter–protomer salt bridges with E462 from the helix alpha1, the secondary structure element found only in severing proteins, and with D559 from PL1. Because mutation of E462 abolishes severing, K555 is important in connecting the ATPase and severing activities of spastin. Importantly, K555 in *D. melanogaster* spastin is the equivalent of K265 in *C. elegans* katanin (Fig. 2E,^31^), which is one of the four crucial positions in katanin with an important role in connecting PL1 with the fishhook. The third special residue, which participates in multiple salt bridges only in the full spastin spiral (ATP+E15) state, is E633 from PL3. It forms inter–protomer salt bridges with R591 and R600, both from PL2, in 3 protomer pairs. The salt bridge E633-R600 is present also in the original cryo–EM structure and the mutation E633A reduces both the ATPase and the severing activity, but does not abolish either. ^29^ Finally, D635 from PL3 participates in 2 intra–protomer salt bridges in the ATP and ATP+E15 states: with R591 and with K603 in most of the protomers. Position D635 is the invariant equivalent of D346 in *C. elegans* katanin and D493 in *H. sapiens* spastin (Fig. 2E^31^). They likely each serve as an anchor between PL3 and oligomerization regions in the respective severing protein.

**Figure 2:**
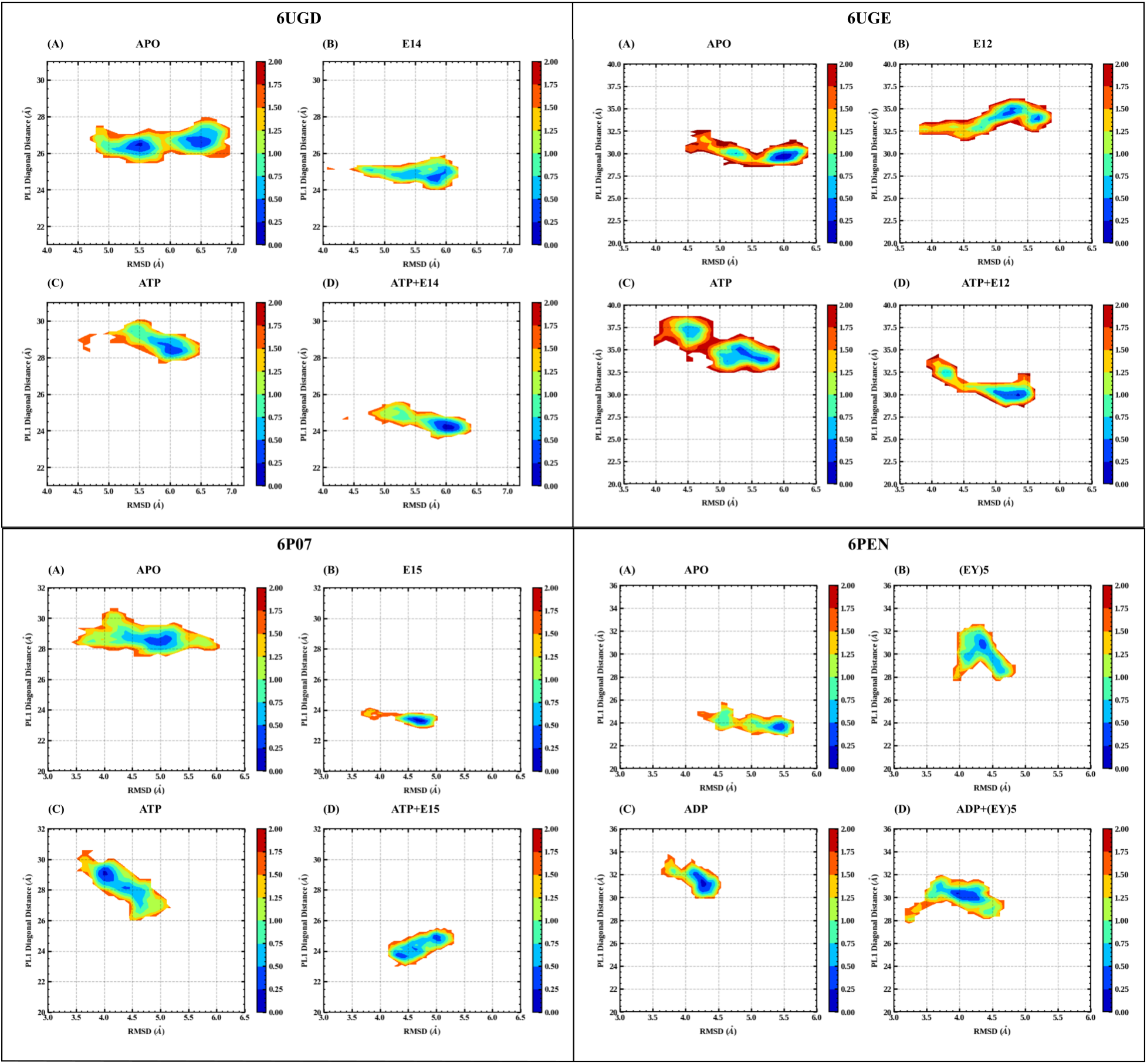
Free Energy Landscape plotted as a function of RMSD and PL1–diagonal distances for the spiral and ring setups of katanin and spastin.

The *H. sapiens* spastin ring is the only state were we found 6 special positions that participate in multiple salt bridges. First, position R450 from PL2, which makes 3 salt bridges in the APO state and 2 salt bridges in the rest of the states. In APO it forms 2 inter–protomer salt bridges with D444 from PL2 and with E491 from PL3 in 3 pairs of neighboring protomers and 1 intra–protomer salt bridge with D493 from PL3 in all 5 protomers that contain ADP in the cryo–EM structure (i.e., in all but protomer F). In the ADP and ADP+EY5 states, R450 makes 2 salt bridges: 1 inter–protomer with E442 from the WB motif in 5 pairs of neighboring protomers or with E491 from PL3 in 3 pairs of neighboring protomers and 1 intra–protomer with D493 from PL3 in all 5 protomers that contain ADP in the cryo–EM structure (i.e., in all but protomer F). Position 450 is the equivalent of R591 in *D. melanogaster* spastin (see Table 2E from^31^). The second position is R460, which makes 2 intra–protomer salt bridges in all states with E420 and with D456 from PL2. The third position is D493 from PL3, which makes 2 intra–protomer salt bridges in all states with R450 from PL2 and with K462. The fourth position is D456 from PL2, which makes 2 salt bridges in all states with the exception of the APO state: 1 inter–protomer with K462 and 1 intra–protomer with R460. The fifth position is E454 from PL2, which makes 2 intra–protomer salt bridges in all states with the exception of the APO state: 1 with R459 and 1 with K462. The sixth and final position is K462, which makes 2 or 3 salt bridges: 2 intra–protomer in the APO state (with E454 from PL2 and with D493 from PL3), and 3 in the other 3 states (1 inter–protomer with D456 from PL2, and the 2 intra–protomer formed in the APO case as well).

In summary, in severing proteins we found networks of salt bridges: (A) centered around position K265 from PL1 in katanin (Fig. S3) and its equivalent K555 position from PL1 in spastin spiral (Fig. S4). It contains 4 members in katanin (D171 from the fishhook element, D269 from PL1 and K272), which form 2 inter- and 1 intra–protomer salt bridges, and 3 members in spastin spiral (E462 from the alpha1 helix, which is special to severing proteins, and D559 from PL1), which form 2 inter–protomer salt bridges. This network connects PL1 loops from neighboring protomers with severing proteins specific functional elements (fishhook and alpha1 helix) involved in the hexamerization. Importantly, this network is absent in the spastin ring; and (B) centered around R301 from PL2 loop in katanin (Fig. S3) and its equivalent R591 position from PL2 in spastin spiral (Fig. S4) and R450 position from PL2 in spastin ring (Fig. S4). This mixed inter- and intra–protomer network contains 4 members in the katanin spiral (E293 from the WB motif, D346 from PL3 and K314), which form 1 inter– and 2 intra–protomer salt bridges, 6 members in the katanin ring (E293 from the WB, D295, E344 and D346 from the PL3 loop and K314) and in the spastin spiral (D585 from the WB motif, E633 and D635 from the PL3 loop, R600 from the PL2 loop, and K603), which form 3 inter– and 2 intra–protomer salt bridges, and 11 members in the spastin ring (D444 from the PL2 loop or the E442 from the WB motif depending on the setup, E491 and D493 from the PL3 loop, K462, E454 and D456 from the PL2 loop, E420, R459, and R460), which form 4 inter- and 6 intra–protomer salt bridges. Whereas network (B) is found in all the states of the severing proteins, we found that it is more extensive in the ring than in the spiral state of an enzyme and that it reaches the maximum size in the spastin ring, where it is centered around position R450 and also that it consists of the maximum number of intra–protomer salt bridges. These findings strongly suggest that the stability of the ring state, especially in spastin, is dependent on its extensive network of positions inside each protomer that can form long-lived complex salt bridges with each other. Moreover, because the protomer lacking the nucleotide in the ring state is excluded from the network, our results show that the formation of the salt bridge networks in severing proteins is dependent on the presence of the nucleotide.

Structural studies highlighted a network of salt bridges formed between, on the one hand, the basic residues K250 and R252 of the canonical pore loops of NBD1 and, on the other, by E254 and E256 in counterclockwise and clockwise neighboring protomers (Fig. S5), respectively, to provide key stabilizing interactions that support translocase and disaggregase activities of the ClpB hexamer. ^25^ The important collective contribution of these salt bridges for the chaperone function was confirmed by point mutations K250E and R252E that result in defective disaggregase activity. ^25^ Prompted by these observations, we probed the dynamics of salt bridges formed in all pore loop regions (PL1-4) of NBD1 and NBD2. Table 1 summarizes the number of salt bridges observed in both ring (6OAX) and spiral (6OAY) states of ClpB and Supplementary Table S6 indicates the specific salt bridge pairs observed with the highest frequency in our simulations. Notably, in both ring and spiral states, the number of inter–protomer salt bridges formed by NBD1 amino acids is larger than that of the intra–protomer salt bridges, which indicates that stabilization of the hexameric structure requires extensive interactions in this flexible region. In the ATP- and substrate–bound state, we observed enhancement of inter–protomer salt bridges in spiral vs. ring configurations indicating the functional role of hexamer stabilization during the ClpB cycle to modulate substrate gripping. In accord with these observations, non-functional ring or spiral configurations obtained through removal of ATP and/or substrate contain similar numbers of salt bridges. We found that specific NBD1 inter–protomer salt bridges K250-E254 and E256-R252, highlighted by Rizo et. al., have strong dynamic contribution to the ClpB hexamer stabilization, with K250-E254 being found in all ClpB states probed and E256-R252 present in the substrate–bound state (ALA) and formed at all inter–protomer interfaces except the seam. The inter–protomer PL1 network also includes E257-K250 salt bridges, which were found at most interfaces in all configurations of both ring and spiral states. Stronger hexamer stabilization of the ATP- and substrate–bound spiral state is effected through inter–protomer salt bridges coupling PL1 and PL2 residues D290-K250, which are absent in the ring state and are replaced by the intra–protomer D290-R252 salt bridge in the ATP– and substrate–bound as well as APO ring configurations, and D290-K288 salt bridges forming an inter–protomer PL2 network in APO, ALA, and ATP ring configuration.

In contrast to the large variability of the number of salt bridges in NBD1, we found nearly invariant numbers of intra- and inter–protomer salt bridges in NBD2 in spiral and ring states, which underscores its stable interaction with the substrate. The most frequent inter–protomer salt bridge in NBD2 is E639-K640, which underlies the network formed by PL4 loops. In addition, the ubiquitous intra–protomer salt bridge involving the E636-R645 pair, present in all states in most of the protomers except the seam, connects the two ends of PL4. Although salt bridge networks involving these residue pairs were not identified in the cryo–EM structural studies, point mutations at positions 639 and 640 were found to result in large reduction of disaggregase activity. ^25^ The distinct pattern of salt bridges in NBD2 compared with NBD1 can be attributed to the absence of charged residues in PL3, which includes the characteristic GYVG loop motif that precludes formation of inter–protomer salt bridge networks or inter–loop salt bridges within the same protomer.

Overall, these findings recapitulate our results for the spiral and ring katanin structures, and provide strong support for the hypothesis that collective pore loop stabilization is required for optimal interaction with the substrate. We surmise that strong inter–protomer salt bridge networks result in formation of PL rings that help stabilize the oligomeric structure of ClpB and enable substrate gripping during the functional cycle. Distinct salt bridge network patterns of the two AAA+ domains highlight their functional specialization into variable substrate gripping during the allosteric cycle by NDB1 and the disaggregase role of NBD2.

### 4.3. Distinct central pore dynamics of severing proteins and ClpB upon nucleotide– and substrate–binding indicates divergent substrate remodeling action

Next, we considered the central pore dynamics of the severing proteins hexamers during the allosteric cycle. To determine the main states populated in our simulations by the functional pore loops in the various hexameric structures as well as their changes driven by the binding of co-factors, we analyzed the free energy landscapes built from the probability density in the (PL1, RMSD) space (Fig. 2) and, respectively, in the (PL2,RMSD) space (Fig. S6 and S7). As shown in Fig. 2, the presence of the substrate in the spiral state, especially when combined with the presence of ATP, has a strong effect on pore dynamics through constraints imposed on PL1 loops that keep the channel in a tight configuration with an average pore diameter (measured as the diagonal distance between PL1 loops that changes the most during our trajectories between the various setups) of approximately 24 Å in both katanin and spastin, compared to an average diameter of 29 Å in the absence of the substrate peptide. This effect is particularly striking in the case of spastin (~26% decrease in pore size in the presence of the substrate peptide) (Fig. 2[6P07]). In all cases and for both severing proteins, the presence of the substrate leads not only to the contraction of the diameter of the central pore but also to a reduction in the degree of variation in the structure as measured by the RMSD (Fig. 2[6UGD] and 2[6P07]). Moreover, we found that in all our simulations with the substrate the tight spiral organization of the pore loops from the 6 protomers around the SP, seen in the original cryo–EM conformation, ^30^ is maintained for the duration of the trajectory (Fig.S10[b,d]). In contrast, the absence of any binding partners (the APO state) or the binding of only the ATP to the spiral state result in a partial loss of the spiral arrangement. For katanin the PL1 loops from the 6 protomers separate into two sets in the ATP trajectories: ABC and DEF. This is due to the movement of protomers C and D away from each other and consequently in the vicinity of their other neighbors in the respective cluster (Fig. S9[c,g]). In spastin the enlargement of the pore in the absence of the substrate is more often the result of chain F the drifting away from the rest of the protomers (Fig.S10[a,c]). In all cases, the APO spiral state has the largest structural variability as measured by the RMSD.

In the ring state we found that the central pore opens up when either the nucleotide alone or the substrate alone is bound to the hexamer: the average PL1 diameter is 36 Å in katanin and ~32 Å in spastin. For katanin this is due to the drifting of chain A (the protomer without ATP) away from the rest of the protomers (Fig. S9[f-g]), whereas in spastin it is the corresponding movement of chain F (the protomer without ADP) away from the other chains (Fig. S10[f-g]). In both severing proteins the pore closes in the absence of any binding partners leading to APO conformations with PL1 diameters of 30 A in katanin and 23.5 Å in spastin (Fig. 2[6UGE] and 2[6PEN]), which is the exact opposite of the behavior from the spiral case. Interestingly, only in the spastin ring the pore opens upon binding of both the nucleotide and the substrate (full spastin ring case) reaching a diameter of 30.5 Å, which, again is the opposite behavior from the spiral case. We also found that, for the katanin ring structure, the binding of the nucleotide (ATP) to 5 of its 6 protomers or the binding of the minimal substrate increases the structural variability, as seen in the increase of the RMSD compared to the APO state (Fig. 2[6UGE]). In the spastin ring, we found the opposite behavior: the binding of either the ADP to 5 of the 6 protomers or the binding of (EY)5 leads to a reduction in the RMSD compared to the APO state (Fig. 2[6PEN]), which recalls the behavior from the spiral case. Thus, in most cases, the binding of the nucleotide to the ring state of a severing protein has the opposite effect on the structural variability of the protein, as measured by the global RMSD, compared to its binding to the spiral state of the same protein.

For the FEL based on the probability density in the (PL2,RMSD) space in the spiral state of the katanin structure (Fig. S6[6UGD]) we found that the diagonal distance between PL2 loops shrinks only minimally (by ~12%) when the substrate peptide (E14) is bound in the pore compared to the no co–factors bound (the APO) state. The presence of the ATP (alone or together with the peptide) leads to a slight ~12% increase in the separation between PL2 loops compared to the APO state. For the spastin spiral case (Fig. S7[6P07]), the SP (either alone or together with ATP) leads to a larger reduction (by ~20%) in the PL2 separation compared to the APO state, whereas binding of ATP alone has no effect on the distance between the diagonal PL2 loops compared to the APO state. Thus, for both katanin and spastin spiral the changes in the diagonal distances between the PL1 and, respectively, the PL2 loops show similar contraction in the presence of the substrate, but the changes are much more pronounced for the PL1 loops. The increase in the separation between PL2 loops in the presence of ATP is bigger in the ring state of katanin (Fig. S6[6UGE]) as it increases by ~25% compared to the full ring case (reaching a diameter of 26 A in the ATP compared to 23 Å. in the ATP+E12 case). For the spastin ring (Fig. S7[6PEN]), binding of the minimal substrate ((EY)5) in the presence or the absence of ADP leads to a very large ~50% increase in the distance between diagonal PL2 loops compared to the APO case (from a ~10 A diameter in the APO state to an average diameter of 15.5 Å in the states with the substrate bound), whereas the presence of the nucleotide alone leads to a ~35% increase compared to the APO case. Again, the changes in the diagonal PL2 loops distances in the ring states of katanin and spastin follow the changes seen in the PL1 loops distances. However, whereas for the katanin ring the changes in the PL2 loops are more modest than those in the PL1 loops, for the spastin ring the PL2 loops are the ones that exhibit the larger changes upon co-factors binding.

The fact that the PL1 and PL2 distances change in a similar manner upon nucleotide or/and SP binding, but the PL1 changes are more pronounced supports a model in which both loops sense the presence of co-factors bound to the spiral state, with the PL1 loops driving the adjustments in the size and shape of the central pore. This is especially true in the case of the spastin spiral, which exhibits the most dramatic changes in the diagonal PL1 distances. In contrast, in the ring state of spastin, which is the only hexameric structure solved in the presence of ADP,^31^ the PL2 loops adjust the most upon binding of both ADP and the substrate resulting in the opening of the central pore. Previously we showed that the spastin ring is the only state characterized by one large network of inter– and intra–protomer salt bridges that connects positions in PL2 with PL3, the nucleotide–binding regions, and positions involved in the oligomerization of the hexamer (Fig. S4[6PEN]). We propose that the changes in the hexameric state upon ADP and substrate binding, which drive the adjustment of the PL2 loops with respect to one another leading to the opening of the pore, are able to propagate to the PL2 loops through the positions that form this salt bridge network. Next, we considered the central pore dynamics of the ClpB hexamer during the allosteric cycle. As in the case of single–ring AAA+ proteins discussed above, we discerned motions that underlie pore dynamics by examining FEL maps in the (RMSD, PL) space of each configuration. To glean characteristic motions on each NBD ring, we focused on their respective canonical loops, PL1 and PL3. As shown in Fig. 3 and S8, the presence of the substrate has a strong effect on pore dynamics in NBD1 through constraints imposed on PL1 loops that maintain the channel in a configuration favorable for substrate gripping and translocation with an average pore diameter of approximately 20 Å in the functional ATP– and substrate–bound spiral structure and a slightly larger diameter of 21 Å in the nucleotide–free and substrate–bound structure (ALA). Consistent with this observation, removal of the substrate (APO and ATP–bound configurations) resulted in a narrower channel width of approximately 18 Å as flexible PL1 loops are able to probe their conformational space more broadly. Substrate–bound configurations also correspond to larger pore width in the ring state, with an average diameter of approximately 22 Å in the functional ATP- and substrate–bound structure (ALA+ATP) and 22.2 A in the nucleotide–free and substrate-bound structure (ALA), whereas substrate–free configurations are 10–14% narrower, with diameters of approximately 19.5 Å (ATP) or 20 Å (APO). We note that spiral structures include narrower pores, consistent with a higher substrate affinity of ClpB and stronger interaction of protomers with the polypeptide at the initiation of the active translocation step. By contrast, larger pore diameter associated with the ring structures is consistent with weaker substrate affinity in this state and release of the polypeptide chain by the ClpB protomer that promotes translocation. In both ring and spiral structures, substrate– and nucleotide– bound conformations are also characterized by reduced conformational flexibility, indicated by smaller backbone RMSD values, compared with APO, ALA and ATP structures. These trends in the size of the pore in the spiral versus the ring state and in the backbone RMSD in the nucleotide and substrate bound states are the same as in the severing proteins. This finding suggests that the central PL1 loops in ClpB and in severing proteins undergo similar conformational changes between the spiral and the ring states upon ATP hydrolysis. However, changes in pore diameter observed in spiral ClpB structures occur in the opposite direction compared to those identified in severing proteins. In the latter case, for example, substrate–binding was found to correspond to a reduction of the pore diameter that enables strong substrate gripping. We propose that strong modulation of hexameric conformations and increased central pore width of ClpB by nucleotide and substrate–binding are consistent with the versatility to accommodate diverse substrates, such as single and multiple polypeptide chains, and the ability to support the double–ring structure through a stable network of inter–protomer interactions. The opposite behavior found in severing proteins points towards a different substrate recognition and processing mechanism: severing proteins likely exhibit high substrate specificity and continuous gripping of substrate, which does not allow for repetitive application of mechanical force to induce unfolding and translocation of the substrate. This recalls our previous findings that a pushing (wedging), rather than an unfoldase, action for severing proteins leads to statistically identical distributions compared to the distributions from in-vitro katanin severing assays.^78,79^

**Figure 3:**
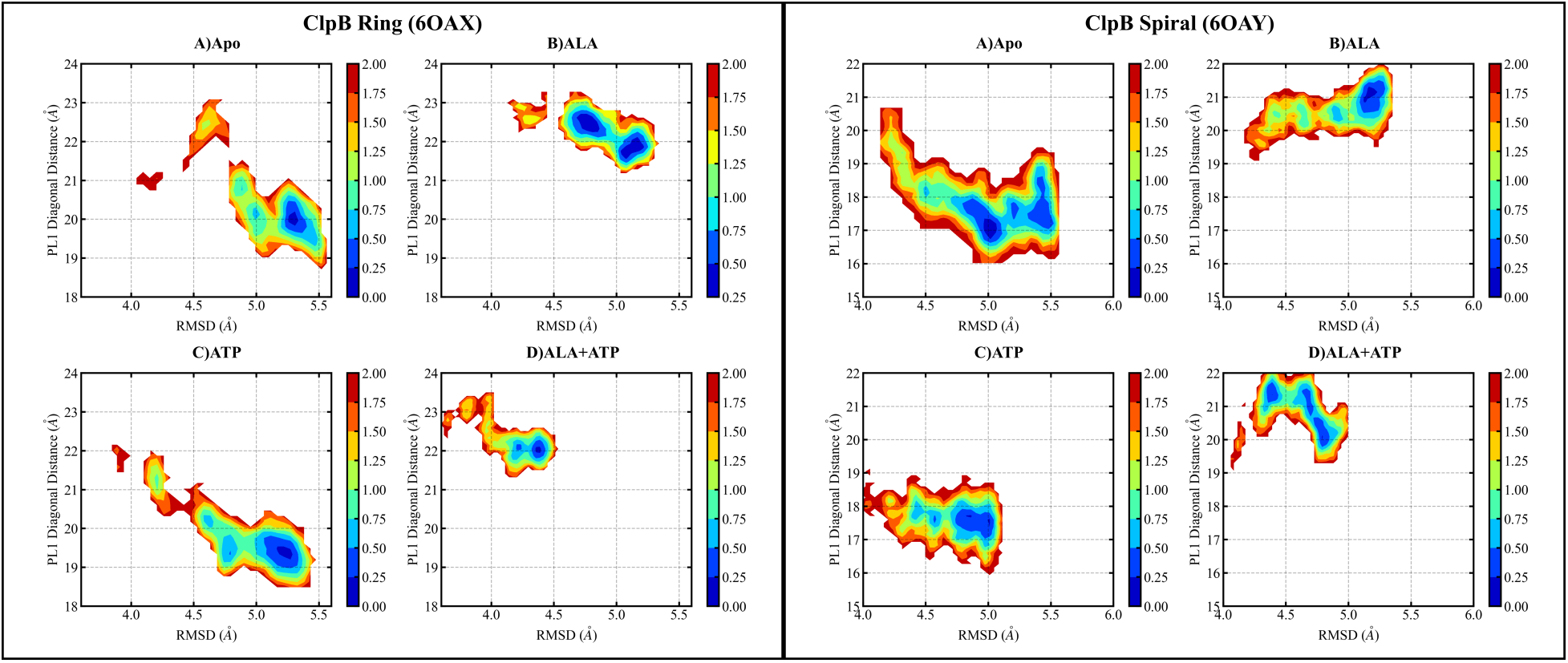
Free Energy Landscape plotted as a function of RMSD and PL1–diagonal distances for ClpB ring and spiral states State: A) ClpB APO state, B) ClpB with substrate bound, C) ClpB with ATP, and D) ClpB with both substrate and ATP bound.

The pore constriction corresponding to the NBD2, which was characterized by monitoring the diagonal distance between PL3 loops, was found to display a similar behavior as the one for NBD1. As shown in Fig. S11, substrate presence in ALA and ALA+ATP structures maintains the pore in the favorable configuration for substrate gripping and translocation, with a diameter of about 22 Å, in both the spiral and ring states. In accord with the NBD1 behavior, removal of the substrate and nucleotide, corresponding to the APO structure, results in narrower NBD2 pore constrictions with average diameter of 19.5 Å. in the spiral configuration and 20 Å in the ring configurations. Nevertheless, in contrast to the NBD1 case, nucleotide–bound and substrate–free configurations of the ring and spiral states display distinct behavior. Whereas in the spiral state the presence of nucleotide results in a narrower pore constriction with an average diameter of approximately 20 Å, in the ring state the pore is wider with an average diameter of approximately 24.5 Å. This divergent behavior is in accord with the different interaction of NBD1 and NBD2 loops with the substrate, namely specific interactions between the charged amino acids of NBD1 loops and non-specific interactions of NBD2 loops.

### 4.4. Divergent intra-ring cooperativity of protomers in severing proteins and of NBD domains of ClpB

We calculated the simplified community networks using the Bio3D package for katanin and spastin hexamers in multiple setups, as described in the Methods (Fig. 4). Strikingly, a common theme is that the majority of clusters from the various setups are intra-rather than inter–protomer. For example, the ring state of spastin (Fig. 4[6PEN]), the only one solved in the presence of ADP, does not show formation of inter–protomer clusters in either of the 4 setups probed in our simulations. In addition, the full (ATP and substrate peptide) spiral and ring states of both katanin and spastin are characterized by the formation of large intra–protomer clusters only. The absence of inter–protomer clusters signals the absence of cooperativity between protomers during severing. However, there are notable exceptions where we found signatures of cooperative action between protomers. In the spiral state of both katanin and spastin the presence of ATP only results in the formation of the largest multi–protomer cluster (> 400 residues) among all the spiral setups (Fig. 4[6UGD] and 4[6P07]). The cluster covers significant portions from the two terminal protomers, A and F: their HBD regions, the ATP binding sites in their N–terminal part, the WA regions and the PL3 loops, as well as most of the NBD region in protomer B. This indicates that the close cooperation between the two terminal protomers and the NBD region of the second protomer is a signature of the spiral state of severing proteins with only ATP bound. Moreover, our results show that when only ATP is bound to the katanin spiral the main regions that cluster together are the ATP–binding sites of the protomers: the ATP–binding regions from the end protomers (A, B, and F) cluster together, whereas the ATP–binding regions from the middle 3 protomers each form an independent cluster. In contrast, in the absence of both ATP and the minimal substrate (E14), the main regions that cluster together in a protomer are those responsible for katanin’s hexamerization: the fishhook positions and the C–terminal region (the alpha11-alpha12 linker, alpha12, and the C–terminal end, i.e., positions 449–472 called the C-Hlx^39^). Another example of cooperativity between protomers is in the spastin spiral state with only the E15 bound, where we found many large inter–protomer clusters (Fig. 4[6P07]). First, that the PL1 loops from 4 protomers (C, D, E, and F) all form one cluster (120 positions), which includes also the PL2 loops from protomers D and E. Second, that the HBD region in protomer A and most of the NBD in protomer B cluster together (209 residues). This is completely different from the behavior of the katanin spiral with E14, where all the clusters are intra–protomer. The commonality between spastin spiral with E15 and katanin spiral with E14 is that most of protomer F forms one large cluster (256 positions) and that the rest of the clusters correspond to the NBD and HBD domains from the various protomers. Thus a strong inter-communication between PL1 loops in adjoining protomers is a characteristic of the spastin spiral in the presence of a minimal substrate peptide. This finding, combined with the results of the free energy landscape analysis in the (PL1,RMSD) space from above, strongly suggest that the large contraction of the PL1 loops in the presence of a substrate for spastin is a local event, being independent of the overall motions of the protomers.

**Figure 4:**
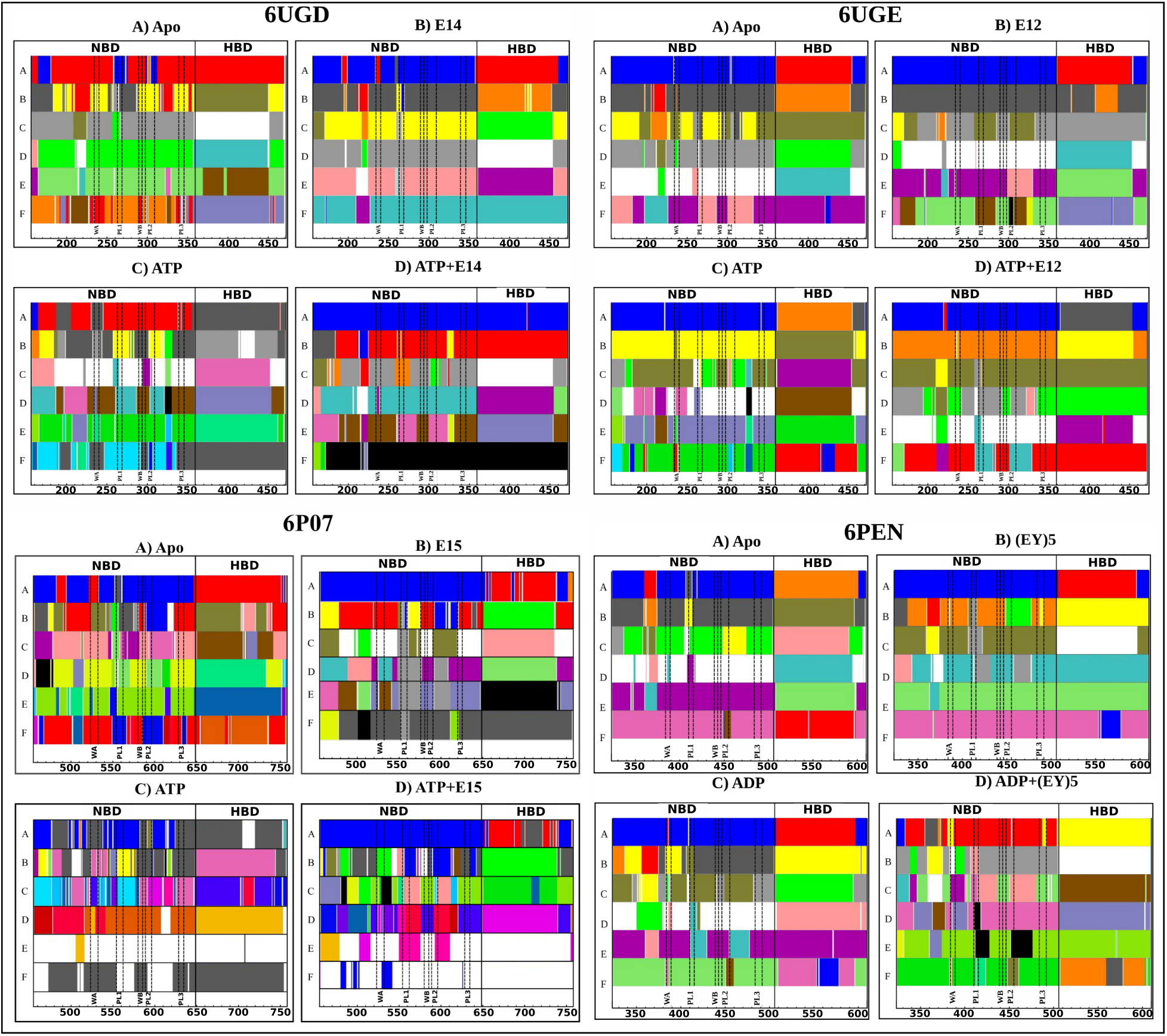
Cluster analysis for katanin and spastin hexamers. a) katanin spiral (6UGD), b) katanin ring (6UGE), c) spastin spiral (6P07), and d) spastin ring (6PEN) showing the NBD, HBD regions and the functional elements, similar to Fig. 1a (WA, WB, PL1, PL2 and PL3).

A third example is in the APO state of the spastin spiral (Fig. 4[6P07]) where we found a large inter–protomer cluster (352 residues) that spans the HBD in protomer A, the NBD and the C–terminal end of the HBD in protomer B, and parts of protomer F (the NBD region, excluding the 3 pore loops, PL1, PL2, PL3, and most of its HBD region). The formation of this cluster is unique to the spastin. The common findings for the spastin and katanin spiral in the APO state are: (i) 5 protomers are divided into clusters that correspond primarily to their 2 structural domains; (ii) whereas the HBD is always found in one cluster, the NBD is split into smaller clusters in all protomers; and (iii) the C-Hlx^39^ clusters together with positions from the NBD in the majority of the protomers (A, B, C, and D). Regarding HBD and ATPase, McNally and collaborators^39^ proposed that HBD undergoes structural changes (unfolding/refolding) as a result of ATP hydrolysis. Our results support this view and provide molecular details of the changes. The RMSF analysis showed that the HBD is the region with the largest fluctuations in all protomers (reaching values that are 2-3 times larger than for any other structural element, up to ~12 A). This finding, combined with the fact that the C–terminal end of the HBD is clustered together with the ATP–binding regions of a protomer, shows that ATP hydrolysis is likely associated with large structural changes (~1 nm) in the HBD region of the spiral state of severing proteins.

Finally, in the ring state of katanin with only ATP bound to all the protomers, with the exception of protomer A, we found a large inter–protomer cluster (242 residues) corresponding to the HBD region in protomer B and most of the NBD from protomer C. The rest of the protomers separate in 2 clusters according to their 2 domains, NBD and HBD. Importantly, this time, unlike in the spiral ATP case, we no longer saw correlations between the nucleotide–binding regions and the HBD domain in the 5 protomers with ATP bound. Due to the importance of ATP hydrolysis to the conformational change in the HBD discussed above, the loss of such strong correlations might explain why in the RMSF plots discussed above the fluctuations seen in the C–terminal end of the HBD are reduced dramatically in the ring versus the spiral state.

As shown in Fig. 5, in each ring and spiral configuration, the calculated simplified community networks of ClpB highlights four distinct regions of protomers, which can be mapped onto the large (L) and small (S) structural domains of the two NBDs, namely NBD1–L (residues 161–340); NBD1–S (residues 341–408); NBD2–L (residues 555–765); and NBD2–S (residues 766–85 3).^80^ The major aspect revealed by clustering patterns of ClpB is coupling of L and S domains within the same NBD of neighboring protomers in the counterclockwise direction. Allosteric networks resulting from coupling within each NBD hexamer, in conjunction with the salt bridge networks noted above, underscore coordinated action of ClpB protomers in their interaction with the substrate. Nevertheless, asymmetric behavior is identified in the two NBD rings, with substrate-independent inter–protomer coupling of NBD1 regions and substrate–induced coupling of NBD2 regions. As shown in Fig. 5, in the spiral configuration, inter–protomer coupling of NBD2 regions is reduced upon removal of the substrate, corresponding to APO and ATP–bound configurations, which supports substrate–induced coordination within this domain, whereas persistent inter–protomer coupling is present within NBD1 in substrate–bound as well as substrate–free configurations. In addition, in the functional ATP– and substrate–bound configuration, multi-protomer clusters formed by PL3 loops of non-seam protomers provide support for the ClpB interaction with the substrate. Consistent with these observations, reduced substrate affinity of ring structures is found in substrate–bound configurations as shown by persistent inter–protomer coupling in NBD1 and weaker coupling in NBD2. In these configurations, inter–protomer clusters in NBD2 are replaced by large (~300 residues) intra–protomer clusters comprising both L and S regions and substrate grip is maintained only through the multi–protomer cluster comprising PL3 loops of ATP–bound protomers. These observations are consistent with the functional role of the NBD2 domains in the translocase action as well as with multi– protomer ClpB interactions with the substrate in structures highlighted by Rizo et al^25^ and in ClpX structures determined by Fei et al.^28^ As noted above, such inter–protomer coupling is not universally observed in single–ring hexameric structures of severing proteins, therefore we surmise that it provides specific functional support for mechanisms of the double–ring structure of ClpB.

**Figure 5:**
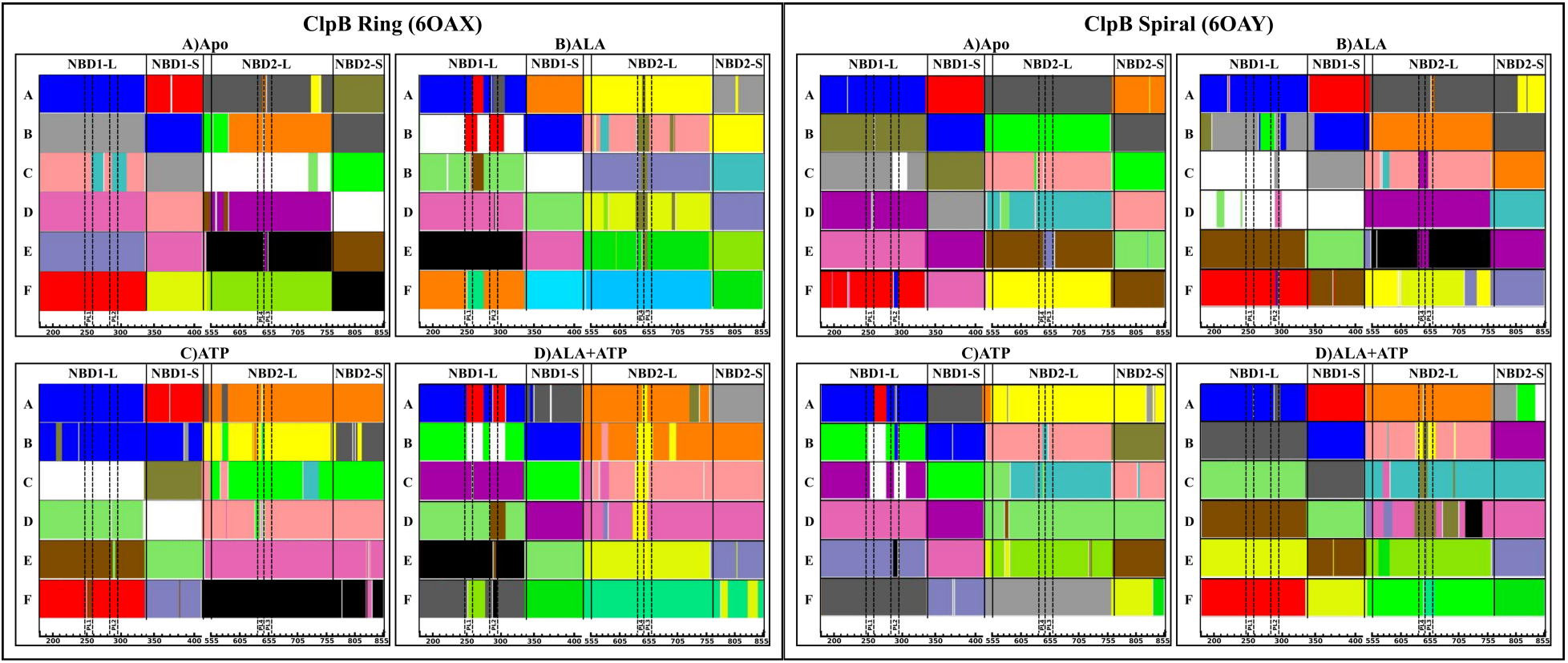
Cluster analysis for ClpB hexamers. a) ClpB ring (6OAX), and b) ClpB spiral (6OAY) showing the NBD1, NBD2 regions and the functional elements, similar to Fig. 1b (PL1, PL2, PL3 and PL4).

### 4.5. Internal motions and relaxation times indicate coupling between collective and local motions

To gain insight into the internal motions responsible for the function of AAA+ motor proteins from our simulations, we determined the timescales for dynamical processes inside the hexamers in the various states. For the severing proteins, we calculated, for each protomer, the autocorrelation function (ACF) of the distance between the C_α_ atom of a central and highly conserved amino acid in PL1 (R267 for katanin, Y556 for spastin spiral, and Y415 for spastin ring) and the center of mass (COM) of the protomer. We fitted the ACF using Eq. 3 (see examples in Fig. S12 from the SI), which is a combination of a stretched exponential and a single exponential.^75^ Next we determined the characteristic relaxation time of the total decay, *τ*^*^, as the integrated area of the autocorrelation function, according to Eq. 4.^75^ We extracted *τ*^*^ from the average (over all the trajectories of a given length) ACF decay in each protomer and in each state from our runs at 5 ns, 25 ns, and 50 ns. While the ACF is calculated for only up to 50% of a trajectory, we found, as seen by others as well,^81,82^ that it becomes negative well before this time point due to the lack of statistics at long times. Thus, for each trajectory length we used the ACF data from only up to the first 20% of the trajectory beyond the initial equilibration time (for example, after the first 5 ns from a 25 ns long trajectory and after the first 1 ns from a 5 ns trajectory). Moreover, for each protomer in each state, unless we could fit the ACF through the entire time interval with high confidence (low χ^2^ value, i.e., < 0.001), we used the dependence of the χ^2^ versus time to select the statistically significant interval for the fit of the ACF by Eq. 3: typically, we selected the time interval corresponding to the elbow in the χ^2^ vs time plot, i.e., we fitted the ACF only up to the time when the χ^2^ exhibits a substantial jump in magnitude (see an example in Fig. S14 for the 6P07 APO setup). The results in Tables S7 and S8 show that for most cases the exponent in the stretched exponential *β* is ~0.5, which agrees with the value of the exponent at 300 K reported in previous work.^75,82^ The physics behind the relaxation time resulting from fitting to a stretched exponential with a sub–unitary exponent (α) corresponds to a purely elastic response preceding a viscous decay. In all setups and for all protomers we found that *τ*^*^ is between T1 and T2 (see Tables S7 and S8), which again recalls the findings from^75^ regarding the behavior of the characteristic relaxation time of the decay in the backbone correlation function for the hen egg white lysozyme. Using the data from the various length trajectories, illustrated in Fig.6 A) and B) for katanin and spastin, respectively, we found that *τ*^*^ = 0.055 * t^1.06^ for katanin and *τ*^*^ = 0.071 * t^0.97^ for spastin, signaling that the decay of the ACF is observation–time–dependent. The power law fit recalls the dependence on the length of the MD simulation trajectories of the characteristic relaxation time of the ACF for distances in monomeric globular proteins, as well as the dependence of the relaxation time on the duration of observation in single molecule FRET experiments reported in the literature.^76,83,84^ The shifting of the ACF towards longer lag times with increasing t, which is a signature of ageing and observation–time–dependent dynamics, has been attributed to a confined subdiffusive continuous time random walk over the protein energy landscape with a superimposed noise. ^76^ Following these studies, we connected our simulation results with the characteristic relaxation times measured in single–molecule FRET experiments ^76,83,85^ by extrapolating the calculated power–law dependence of *τ*^*^ on the length (t) of the measurement leading to a predicted value for *τ*^*^, at the typical experimental timescales from FRET experiments (t =1 ms), of 139 μs for katanin and 50 μs for spastin. The individual values for the *τ*^*^ fitting and the respective predicted *τ*^*^ values at 1 ms observation time for each of the setups in katanin and spastin are in Table S11. This table shows that the predicted values range between a couple microseconds to hundreds of microseconds. Common among all the katanin and spastin structures is that the binding of the nucleotide (ATP or ADP) alone results in the fastest decay of the ACF. In contrast, the ACF in the absence of any binding partners (the APO setup) exhibits the longest decay among the ACFs for all the setups probed in the severing proteins. In order to study the internal motions occurring during conformational changes in the ClpB nanomachine in various states (APO, ALA, ATP and ALA+ATP), we calculated, for each protomer, the ACF of the distance between the C_α_ atom of a central and highly conserved amino acid in PL1 (R252) and the center of mass (COM) of the protomer. The results shown in Tables S9 and S10 show that the average exponent in the stretched exponential is ~0.6, with individual values ranging from 0.4 to 0.7, which is only slightly higher than the corresponding number in severing proteins. Similar to our findings in severing enzymes, in all setups and for all protomers, *τ*^*^ lies between the T1 and T2 values (see Tables S9 and S10). Using the data from the various length trajectories, illustrated in Fig. S15 and reported in Table S12 for all the different setups in ClpB, we found that *τ*^*^ = 0.102 *\** t^0.91^, consistent with our above finding for severing proteins, which indicates that the decay of the ACF is observation–time–dependent. Extrapolation to the typical experimental timescale from FRET experiments of 1 ms leads to a predicted value for *τ*^*^ of 30 μs. This is smaller than the predicted values from the severing proteins. In general, we found that the *τ*^*^ values obtained for the ClpB states (which are up to 110 μs) are shorter than the *τ*^*^ values for katanin and spastin (which are up to 370 μs). Another difference between ClpB and severing proteins is that the slowest decay in the ACF for ClpB corresponds to the state when both the nucleotide and the substrate are bound to the hexamer, whereas the fastest decay corresponds to the APO state. Very recent smFRET experiments,^67^ carried out over one millisecond time windows, investigated the pore–loop dynamics in ClpB and extracted characteristic relaxation times for PL1 motions. The combined plot of these values together with those from our simulations in Fig.6 C) yields the power–law dependence *τ*^*^ = 0.087 * t^0.95^, indicating excellent agreement between experiments and simulations. The resulting *τ*^*^ value on the 1 ms timescale is 43 μs, which agrees with the above computational prediction.

**Figure 6:**
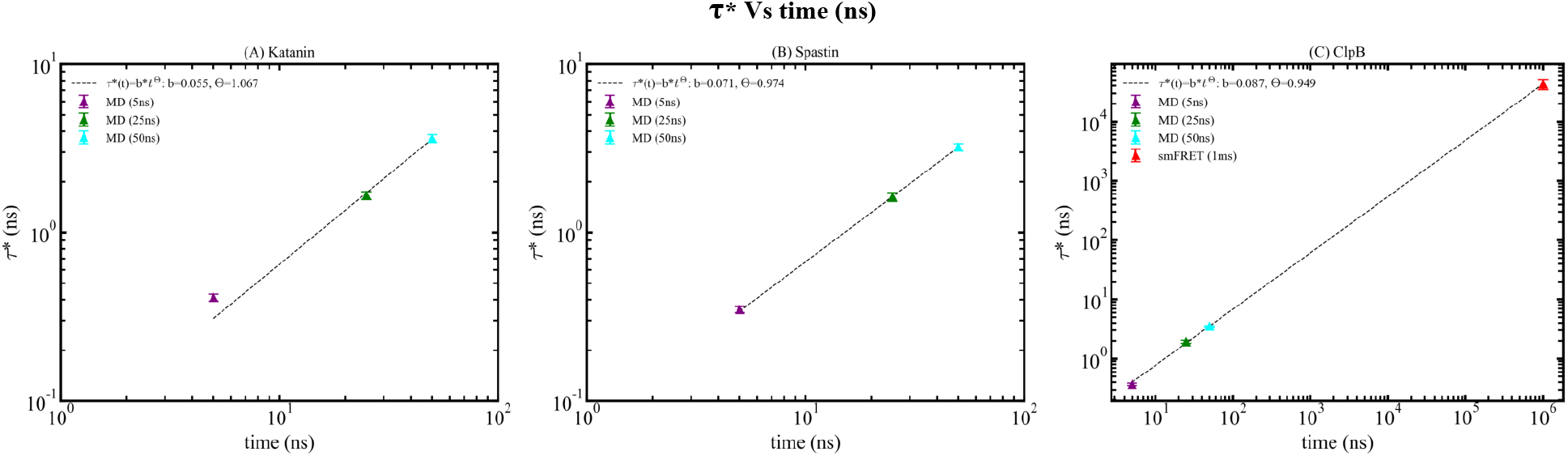
Dependence of relaxation time *τ*^*^ on the observation timescale. Log–log plots indicate power law dependence (dashed line) of *τ*^*^ vs time (ns) for A) katanin (MD data) and B) spastin (MD data) and C) ClpB (MD + smFRET data). Data from MD simulations of 5 ns (magenta), 25 ns (green) and 50 ns (cyan) trajectories, as well as from smFRET experiments (red)^67^ are shown. Standard error of *τ*^*^ data is shown as error bars.

In summary, our result, for both classes of AAA+ proteins, that the characteristic ACF decay time *τ*^*^) lies between T1 and T2 indicates that in these proteins there is substantial coupling between collective and local motions in the hexamers. ^75^ This coupling aligns well with our above reported formation of networks of intra– and inter–protomer complex salt bridges that help organize the main functional regions of the hexamers: the central pore loops, the ATP–binding sites, and the hexamerization sites. Moreover, both classes of proteins exhibit pronounced ageing and observation–time–dependent dynamics as seen in the shifting of the ACF towards longer lag times with increasing t, ^76^ resulting in a predicted value for *τ*^*^ in ClpB of tens of microseconds on the typical experimental timescale from FRET experiments, which matches very well the experimental value. ^67^ Interestingly the respective predicted value in spastin matches the result for ClpB, whereas in katanin it is longer than in ClpB, which suggests that katanin exhibits slower PL1 kinetics.

## 5. Conclusion

Results of computational studies presented in this paper reveal factors that dynamically stabilize non–planar hexameric structures of AAA+ machines involved in disaggregation or microtubule–severing and underlie conformational dynamics that enables their functions. In all the hexameric states of the severing proteins we found networks of amino acids from the 3 pore loops involved in the formation of long lived complex salt bridges (between at least three charged residues) with each other. Importantly, these salt bridges are not present in the original cryo–EM structures of the severing protein, but rather form during the MD simulations and are thus present in many of the conformations sampled during the runs. Some of these complex salt bridges are inter–protomer, recalling findings from the literature that cooperative networks of charged residues are more often found at subunit–subunit interfaces. ^86^ In proteins, such as in the lipase family, the presence of networks of electrostatic interactions across protein–protein interfaces stabilizes the geometry of the active site and results in specificity in the formation of the complex with its ligands. ^73^ Importantly, while in general there are no rules that state that a salt bridge has a stabilizing effect on a protein structure, networked salt bridges are stabilizing. ^73^ Based on these findings, it is likely that the networks of salt bridges involving the central pore loops in severing proteins play a stabilizing role for the structures of the hexameric assemblies. Furthermore, they are at least in part responsible for the specificity of the formation of the complex with substrate peptides. Finally, the fact that the network that connects PL2 with PL3, which is the most extensive network of electrostatic interactions in severing proteins, shows changes in the identity of the salt bridges depending on the number and types of their binding partners is indicative of the adaptation of this network to enable the recognition of several ligands by a single molecule. ^73^ Based on the salt bridge analysis of our simulations we also propose that, whereas electrostatic interactions are involved in stabilizing the central pore loops, the formation of these contacts is dependent on the presence of the nucleotide and it is driven by different processes depending on the state of the severing proteins: in katanin and spiral state of spastin the structural flexibility of the full hexamer is responsible for the interactions as the majority of the salt bridges are inter protomer, whereas in spastin ring internal motions in protomers lead to the interactions as this state is characterized by a very large number of intra–protomer salt bridges. This proposal is supported by the results of our cluster analysis, which show that correlated motions of residues in hexamers lead to the formation of inter– protomer clusters for both katanin and the spiral state of spastin, whereas the ring state of spastin is characterized by intra–protomer clusters. These findings are mirrored by our results for ClpB, particularly for the NBD1, where the networks of salt bridges that stabilize the arrangement of the central pore loops are primarily inter–protomer and for which the clustering analysis revealed the formation of a large number of inter–protomer clusters. We thus propose that the formation of networks of complex inter–protomer salt bridges, which is dependent on the presence of the nucleotide, and it is driven by the correlated motions of many regions in the hexameric structures, is responsible for the stability of the central pore loops, which is required for functional reasons in AAA+ machines. Furthermore, the adaptation of such networks enables the recognition of several ligands by a given molecular machine. Specialized roles of the two NBD domains of ClpB are highlighted by the distinct pattern of salt-bridge network and inter–protomer coupling. Whereas the NBD1 ring involves cross–protomer salt bridges between residues of PL1 loops and coupled motions of large and small domains of neighboring protomers, the NBD2 ring lacks a salt bridge network stabilizing PL3 loops and includes intra–protomer coupling of large and small domains. These results are consistent with findings of biochemical studies that indicate cooperative interactions of the NBD1 protomers and probabilistic interactions of the NBD2 protomers. ^56^

The presence of substantial correlations between residue motions for the duration of the simulations is also supported by the results of our analysis of the relaxation dynamics of the ACF for the distance between PL1 and the center of mass of the protomer. Our results show that for both severing proteins and ClpB we are in the regime where there is substantial coupling between collective and local motions. We also found that the characteristic relaxation time for the ACF in the hexameric states experiences ageing and observation–time– dependent dynamics, which is the result of a confined subdiffusive continuous time random walk over the protein energy landscape with a superimposed noise, similar to the findings for monomeric globular proteins from the literature. ^76^ Using the resulting power law dependence, we predict that the typical ACF relaxation time is ~45 μs on the timescale of single–molecule FRET experiments, which are carried out for a duration of ~1 ms. Timescales corresponding to these relaxation times are in excellent accord with those identified in very recent smFRET experiments for functional motions of the ClpB nanomachine. ^67^ This finding indicates that, despite the inherent timescale limitations of MD simulations, the motions observed in our work are relevant over many timescale orders, as the dynamics is non-equilibrium and self-similar at least up to the duration of smFRET experiments.

## Author Contributions

R.I.D. and G.S. designed the research. M.D., A.D., R.A.V. carried out all simulations, M.D., A.D., R.A.V., G.S. and R.I.D. analyzed the data and wrote the article.

## Acknowledgement

The authors thank Sue Wickner, Shannon Doyle, Gilad Haran and Hisham Mazal for useful discussions. This work was supported by grants from the National Science Foundation: MCB–18179488 (to RID), and MCB–1516918 (to GS). The simulations were performed using Ohio Supercomputer Center (OSC) resources through project PES0791 (to RID) and Extreme Science and Engineering Discovery Environment (XSEDE) resources, which are supported by the NSF (grant no. ACI–1548562), using the Bridges platform at the Pittsburgh Supercomputer Center through projects TG–MCB200201 (to RID) and TG–MCB170020 (to GS).

## Supporting Information Available

An online supplement to this article can be found by visiting bioRχiv Online at https://www.biorxiv.org.

## References

(1) Glover, J. R.; Lindquist, S. Hsp104, Hsp70, and Hsp40: a novel chaperone system that rescues previously aggregated proteins. Cell 1998, 94, 73–82.

(2) Erzberger, J. P.; Berger, J. M. Evolutionary relationships and structural mechanisms of AAA+ proteins. Annu. Rev. Biophys. Biomol. Struct. 2006, 35, 93–114.

(3) Doyle, S. M.; Wickner, S. Hsp104 and ClpB: protein disaggregating machines. Trends Biochem. Sci. 2009, 34, 40–48.

(4) Martin, A.; Baker, T. A.; Sauer, R. T. Pore loops of the AAA+ ClpX machine grip substrates to drive translocation and unfolding. Nat. Struct. Mol. Biol. 2008, 15, 1147.

(5) Lee, S.; Sowa, M. E.; Choi, J.-M.; Tsai, F. T. The ClpB/Hsp104 molecular chaperone—a protein disaggregating machine. J. Struct. Biol. 2004, 146, 99–105.

(6) Roll-Mecak, A.; Vale, R. D. Structural basis of microtubule severing by the hereditary spastic paraplegia protein spastin. Nature 2008, 451, 363–367.

(7) Heuck, A.; Schitter-Sollner, S.; Suskiewicz, M. J.; Kurzbauer, R.; Kley, J.; Schleiffer, A.; Rombaut, P.; Herzog, F.; Clausen, T. Structural basis for the disaggregase activity and regulation of Hsp104. Elife 2016, 5, e21516.

(8) Deville, C.; Carroni, M.; Franke, K. B.; Topf, M.; Bukau, B.; Mogk, A.; Saibil, H. R. Structural pathway of regulated substrate transfer and threading through an Hsp100 disaggregase. Sci. Adv. 2017, 3, e1701726.

(9) Lee, S.; Roh, S. H.; Lee, J.; Sung, N.; Liu, J.; Tsai, F. T. Cryo-EM structures of the Hsp104 protein disaggregase captured in the ATP conformation. Cell reports 2019, 26, 29–36.

(10) Michalska, K.; Zhang, K.; March, Z. M.; Hatzos-Skintges, C.; Pintilie, G.; Bigelow, L.; Castellano, L. M.; Miles, L. J.; Jackrel, M. E.; Chuang, E., et al. Structure of Calcarisporiella thermophila Hsp104 disaggregase that antagonizes diverse proteotoxic misfolding events. Structure 2019, 27, 449–463.

(11) Puchades, C.; Rampello, A. J.; Shin, M.; Giuliano, C. J.; Wiseman, R. L.; Glynn, S. E.; Lander, G. C. Structure of the mitochondrial inner membrane AAA+ protease YME1 gives insight into substrate processing. Science 2017, 358, 1–10.

(12) Andres, H.; Goodall, E. A.; Gates, S. N.; Lander, G. C.; Martin, A. Substrate-engaged 26S proteasome structures reveal mechanisms for ATP-hydrolysis–driven translocation. Science 2018, 362, 1–9.

(13) Dong, Y.; Zhang, S.; Wu, Z.; Li, X.; Wang, W. L.; Zhu, Y.; Stoilova-McPhie, S.; Lu, Y.; Finley, D.; Mao, Y. Cryo-EM structures and dynamics of substrate-engaged human 26S proteasome. Nature 2019, 565, 49–55.

(14) Majumder, P.; Rudack, T.; Beck, F.; Danev, R.; Pfeifer, G.; Nagy, I.; Baumeister, W. Cryo-EM structures of the archaeal PAN-proteasome reveal an around-the-ring ATPase cycle. Proc. Natl. Acad. Sci. USA. 2019, 116, 534–539.

(15) Ripstein, Z. A.; Huang, R.; Augustyniak, R.; Kay, L. E.; Rubinstein, J. L. Structure of a AAA+ unfoldase in the process of unfolding substrate. Elife 2017, 6, e25754.

(16) Gates, S. N.; Yokom, A. L.; Lin, J.; Jackrel, M. E.; Rizo, A. N.; Kendsersky, N. M.; Buell, C. E.; Sweeny, E. A.; Mack, K. L.; Chuang, E., et al. Ratchet-like polypeptide translocation mechanism of the AAA+ disaggregase Hsp104. Science 2017, 357, 273–279.

(17) Zehr, E.; Szyk, A.; Piszczek, G.; Szczesna, E.; Zuo, X.; Roll-Mecak, A. Katanin spiral and ring structures shed light on power stroke for microtubule severing. Nat. Struct. Mol. Biol. 2017, 24, 717–725.

(18) Han, H.; Monroe, N.; Sundquist, W. I.; Shen, P. S.; Hill, C. P. The AAA ATPase Vps4 binds ESCRT-III substrates through a repeating array of dipeptide-binding pockets. Elife 2017, 6, e31324.

(19) Monroe, N.; Han, H.; Shen, P. S.; Sundquist, W. I.; Hill, C. P. Structural basis of protein translocation by the Vps4-Vta1 AAA ATPase. Elife 2017, 6, e24487.

(20) Su, M.; Guo, E. Z.; Ding, X.; Li, Y.; Tarrasch, J. T.; Brooks, C. L.; Xu, Z.; Skinio-tis, G. Mechanism of Vps4 hexamer function revealed by cryo-EM. Sci. Adv. 2017, 3, e1700325.

(21) Sun, S.; Li, L.; Yang, F.; Wang, X.; Fan, F.; Yang, M.; Chen, C.; Li, X.; Wang, H.-W.; Sui, S.-F. Cryo-EM structures of the ATP-bound Vps4 E233Q hexamer and its complex with Vta1 at near-atomic resolution. Nat. Commun. 2017, 8, 1–13.

(22) Yu, H.; Lupoli, T. J.; Kovach, A.; Meng, X.; Zhao, G.; Nathan, C. F.; Li, H. ATP hydrolysis-coupled peptide translocation mechanism of Mycobacterium tuberculosis ClpB. Proc. Natl. Acad. Sci. USA. 2018, 115, E9560–E9569.

(23) White, K. I.; Zhao, M.; Choi, U. B.; Pfuetzner, R. A.; Brunger, A. T. Structural principles of SNARE complex recognition by the AAA+ protein NSF. Elife 2018, 7, e38888.

(24) Cooney, I.; Han, H.; Stewart, M. G.; Carson, R. H.; Hansen, D. T.; Iwasa, J. H.; Price, J. C.; Hill, C. P.; Shen, P. S. Structure of the Cdc48 segregase in the act of unfolding an authentic substrate. Science 2019, 365, 502–505.

(25) Rizo, A. N.; Lin, J.; Gates, S. N.; Tse, E.; Bart, S. M.; Castellano, L. M.; DiMaio, F.; Shorter, J.; Southworth, D. R. Structural basis for substrate gripping and translocation by the ClpB AAA+ disaggregase. Nat. Commun. 2019, 10, 1–12.

(26) Shin, M.; Asmita, A.; Puchades, C.; Adjei, E.; Wiseman, R. L.; Karzai, A. W.; Lander, G. C. Distinct structural features of the lon protease drive conserved Hand-overHand substrate translocation. BioRxiv 2019, 617159.

(27) Twomey, E. C.; Ji, Z.; Wales, T. E.; Bodnar, N. O.; Ficarro, S. B.; Marto, J. A.; Engen, J. R.; Rapoport, T. A. Substrate processing by the Cdc48 ATPase complex is initiated by ubiquitin unfolding. Science 2019, 365, eaax1033.

(28) Fei, X.; Bell, T. A.; Jenni, S.; Stinson, B. M.; Baker, T. A.; Harrison, S. C.; Sauer, R. T. Structures of the ATP-fueled ClpXP proteolytic machine bound to protein substrate. Elife 2020, 9, e52774.

(29) Sandate, C. R.; Szyk, A.; Zehr, E. A.; Lander, G. C.; Roll-Mecak, A. An allosteric network in spastin couples multiple activities required for microtubule severing. Nat. Struct. Mol. Biol. 2019, 26, 671–678.

(30) Roll-Mecak, A. The tubulin code in microtubule dynamics and information encoding. Dev. Cell 2020, 54, 7–20.

(31) Han, H.; Schubert, H. L.; McCullough, J.; Monroe, N.; Purdy, M. D.; Yeager, M.; Sundquist, W. I.; Hill, C. P. Structure of spastin bound to a glutamate-rich peptide implies a hand-over-hand mechanism of substrate translocation. J. Biol. Chem. 2020, 295, 435–443.

(32) Yokom, A. L.; Gates, S. N.; Jackrel, M. E.; Mack, K. L.; Su, M.; Shorter, J.; Southworth, D. R. Spiral architecture of the Hsp104 disaggregase reveals the basis for polypeptide translocation. Nat. Struct. Mol. Biol. 2016, 23, 830–837.

(33) Martin, A.; Baker, T. A.; Sauer, R. T. Rebuilt AAA+ motors reveal operating principles for ATP-fuelled machines. Nature 2005, 437, 1115–1120.

(34) Hersch, G. L.; Burton, R. E.; Bolon, D. N.; Baker, T. A.; Sauer, R. T. Asymmetric interactions of ATP with the AAA+ ClpX6 unfoldase: allosteric control of a protein machine. Cell 2005, 121, 1017–1027.

(35) Aubin-Tam, M.-E.; Olivares, A. O.; Sauer, R. T.; Baker, T. A.; Lang, M. J. Singlemolecule protein unfolding and translocation by an ATP-fueled proteolytic machine. Cell 2011, 145, 257–267.

(36) Cordova, J. C.; Olivares, A. O.; Shin, Y.; Stinson, B. M.; Calmat, S.; Schmitz, K. R.; Aubin-Tam, M.-E.; Baker, T. A.; Lang, M. J.; Sauer, R. T. Stochastic but highly coordinated protein unfolding and translocation by the ClpXP proteolytic machine. Cell 2014, 158, 647–658.

(37) Olivares, A. O.; Nager, A. R.; Iosefson, O.; Sauer, R. T.; Baker, T. A. Mechanochemical basis of protein degradation by a double-ring AAA+ machine. Nat. Struct. Mol. Biol. 2014, 21, 871–875.

(38) Uchihashi, T.; Watanabe, Y.-h.; Nakazaki, Y.; Yamasaki, T.; Watanabe, H.; Maruno, T.; Ishii, K.; Uchiyama, S.; Song, C.; Murata, K., et al. Dynamic structural states of ClpB involved in its disaggregation function. Nat. Commun. 2018, 9, 1–12.

(39) McNally, F. J.; Roll-Mecak, A. Microtubule-severing enzymes: From cellular functions to molecular mechanism. J. Cell Biol. 2018, 217, 4057–4069.

(40) Goloubinoff, P.; Mogk, A.; Zvi, A. P. B.; Tomoyasu, T.; Bukau, B. Sequential mechanism of solubilization and refolding of stable protein aggregates by a bichaperone network. Proc. Natl. Acad. Sci. USA. 1999, 96, 13732–13737.

(41) Haslberger, T.; Zdanowicz, A.; Brand, I.; Kirstein, J.; Turgay, K.; Mogk, A.; Bukau, B. Protein disaggregation by the AAA+ chaperone ClpB involves partial threading of looped polypeptide segments. Nat. Struct. Mol. Biol. 2008, 15, 641–650.

(42) McNally, F. J.; Vale, R. D. Identification of katanin, an ATPase that severs and disassembles stable microtubules. Cell 1993, 75, 419–429.

(43) Bailey, M. E.; Jiang, N.; Dima, R. I.; Ross, J. L. Invited review: Microtubule severing enzymes couple atpase activity with tubulin GTPase spring loading. Biopolymers 2016, 105, 547–556.

(44) Barsegov, V.; Ross, J. L.; Dima, R. I. Dynamics of microtubules: highlights of recent computational and experimental investigations. J. Phys.: Condens. Matter 2017, 29, 433003.

(45) Vemu, A.; Szczesna, E.; Zehr, E. A.; Spector, J. O.; Grigorieff, N.; Deaconescu, A. M.; Roll-Mecak, A. Severing enzymes amplify microtubule arrays through lattice GTP-tubulin incorporation. Science 2018, 361, 1–12.

(46) Diaz-Valencia, J. D.; Bailey, M.; Ross, J. L. Methods in cell biology; Elsevier, 2013; Vol. 115; pp 191–213.

(47) Bailey, M. E.; Sackett, D. L.; Ross, J. L. Katanin severing and binding microtubules are inhibited by tubulin carboxy tails. Biophys. J. 2015, 109, 2546–2561.

(48) Lacroix, B.; Van Dijk, J.; Gold, N. D.; Guizetti, J.; Aldrian-Herrada, G.; Rogowski, K.; Gerlich, D. W.; Janke, C. Tubulin polyglutamylation stimulates spastin-mediated microtubule severing. J. Cell Biol. 2010, 189, 945–954.

(49) Zolkiewski, M. ClpB Cooperates with DnaK, DnaJ, and GrpE in Suppressing Protein Aggregation. J. Biol. Chem. 1999, 274, 28083–28086.

(50) Schlieker, C.; Weibezahn, J.; Patzelt, H.; Tessarz, P.; Strub, C.; Zeth, K.; Erbse, A.; Schneider-Mergener, J.; Chin, J. W.; Schultz, P. G.; Bukau, B.; Mogk, A. Substrate recognition by the AAA+ chaperone ClpB. Nat. Struct. Mol. Biol. 2004, 11, 607–615.

(51) Doyle, S. M.; Hoskins, J. R.; Wickner, S. Collaboration between the ClpB AAA+ remodeling protein and the DnaK chaperone system. Proc. Natl. Acad. Sci. USA. 2007, 104, 11138–11144.

(52) Doyle, S. M.; Shastry, S.; Kravats, A. N.; Shih, Y.-H.; Miot, M.; Hoskins, J. R.; Stan, G.; Wickner, S. Interplay between E. coli DnaK, ClpB and GrpE during Protein Disaggregation. J. Mol. Biol. 2015, 427, 312–327.

(53) Weibezahn, J.; Tessarz, P.; Schlieker, C.; Zahn, R.; Maglica, Z.; Lee, S.; Zentgraf, H.; Weber-Ban, E. U.; Dougan, D. A.; Tsai, F. T.; Mogk, A.; Bukau, B. Thermotolerance Requires Refolding of Aggregated Proteins by Substrate Translocation through the Central Pore of ClpB. Cell 2004, 119, 653–665.

(54) Nagy, M.; Guenther, I.; Akoyev, V.; Barnett, M. E.; Zavodszky, M. I.; Kedzierska-Mieszkowska, S.; Zolkiewski, M. Synergistic Cooperation between Two ClpB Isoforms in Aggregate Reactivation. J. Mol. Biol. 2010, 396, 697–707.

(55) Doyle, S. M.; Shorter, J.; Zolkiewski, M.; Hoskins, J. R.; Lindquist, S.; Wickner, S. Asymmetric deceleration of ClpB or Hsp104 ATPase activity unleashes protein-remodeling activity. Nat. Struct. Mol. Biol. 2007, 14, 114–122.

(56) Doyle, S. M.; Hoskins, J. R.; Wickner, S. DnaK Chaperone-dependent Disaggregation by Caseinolytic Peptidase B (ClpB) Mutants Reveals Functional Overlap in the N-terminal Domain and Nucleotide-binding Domain-1 Pore Tyrosine. J. Biol. Chem. 2012, 287, 28470–28479.

(57) Huang, L.; Kirmizialtin, S.; Makarov, D. E. Computer simulations of the translocation and unfolding of a protein pulled mechanically through a pore. J. Chem. Phys. 2005, 123, 124903.

(58) West, D. K.; Brockwell, D. J.; Olmsted, P. D.; Radford, S. E.; Paci, E. Mechanical Resistance of Proteins Explained Using Simple Molecular Models. Biophys. J. 2006, 90, 287–297.

(59) Wojciechowski, M.; Szymczak, P.; Carrion-Vazquez, M.; Cieplak, M. Protein Unfolding by Biological Unfoldases: Insights from Modeling. Biophys. J. 2014, 107, 1661–1668.

(60) Wojciechowski, M.; Gomez-Sicilia,; Carrión-Vázquez, M.; Cieplak, M. Unfolding knots by proteasome-like systems: simulations of the behaviour of folded and neurotoxic proteins. Mol. Biosyst. 2016, 12, 2700–2712.

(61) Kravats, A. N.; Tonddast-Navaei, S.; Stan, G. Coarse-Grained Simulations of Topology-Dependent Mechanisms of Protein Unfolding and Translocation Mediated by ClpY ATPase Nanomachines. PLoS Comput. Biol. 2016, 12, e1004675.

(62) Javidialesaadi, A.; Stan, G. Asymmetric Conformational Transitions in AAA+ Biological Nanomachines Modulate Direction-Dependent Substrate Protein Unfolding Mechanisms. J. Phys. Chem. B 2017, 121, 7108–7121, PMID: 28675036.

(63) Javidialesaadi, A.; Flournoy, S. M.; Stan, G. Role of Diffusion in Unfolding and Translocation of Multidomain Titin I27 Substrates by a Clp ATPase Nanomachine. J. Phys. Chem. B 2019, 123, 2623–2635.

(64) Sen, M.; Maillard, R.; Nyquist, K.; Rodriguez-Aliaga, P.; Pressé, S.; Martin, A.; Bustamante, C. The ClpXP Protease Unfolds Substrates Using a Constant Rate of Pulling but Different Gears. Cell 2013, 155, 636–646.

(65) Avestan, M. S.; Javidi, A.; Ganote, L. P.; Brown, J. M.; Stan, G. Kinetic effects in directional proteasomal degradation of the green fluorescent protein. J. Chem. Phys. 2020, 153, 105101.

(66) Mazal, H.; Iljina, M.; Barak, Y.; Elad, N.; Rosenzweig, R.; Goloubinoff, P.; Riven, I.; Haran, G. Tunable microsecond dynamics of an allosteric switch regulate the activity of a AAA+ disaggregation machine. Nat. Commun. 2019, 10, 1438.

(67) Mazal, H. A.; Iljina, M.; Riven, I.; Haran, G. Ultrafast Brownian-ratchet mechanism for protein translocation by a AAA+ machine. bioRxiv doi: https://doi.org/10.1101/2020.11.19.3843132020,

(68) Berman, H.; Henrick, K.; Nakamura, H. Announcing the worldwide Protein Data Bank. Nat. Struct. Mol. Biol. 2003, 10, 980–980.

(69) Spoel, D. V. D.; Lindahl, E.; Hess, B.; Groenhof, G.; Mark, A. E.; Berendsen, H. J. C. GROMACS: Fast, flexible, and free. J. Comput. Chem. 2005, 26, 1701–1718.

(70) Schmid, N.; Eichenberger, A. P.; Choutko, A.; Riniker, S.; Winger, M.; Mark, A. E.; van Gunsteren, W. F. Definition and testing of the GROMOS force-field versions 54A7 and 54B7. Eur. Biophys. J. 2011, 40, 843–856.

(71) Skjærven, L.; Yao, X.-Q.; Scarabelli, G.; Grant, B. J. Integrating protein structural dynamics and evolutionary analysis with Bio3D. BMC Bioinf. 2014, 15, 1–11.

(72) Girvan, M.; Newman, M. E. J. Community structure in social and biological networks. Proc. Natl. Acad. Sci. USA. 2002, 99, 7821–7826.

(73) Kumar, S.; Nussinov, R. Relationship between Ion Pair Geometries and Electrostatic Strengths in Proteins. Biophys. J. 2002, 83, 1595–1612.

(74) Humphrey, W.; Dalke, A.; Schulten, K. VMD: Visual molecular dynamics. J. Mol. Graphics 1996, 14, 33–38.

(75) Okan, O. B.; Atilgan, A. R.; Atilgan, C. Nanosecond Motions in Proteins Impose Bounds on the Timescale Distributions of Local Dynamics. Biophys. J. 2009, 97, 2080–2088.

(76) Hu, X.; Hong, L.; Dean Smith, M.; Neusius, T.; Cheng, X.; Smith, J. C. The dynamics of single protein molecules is non-equilibrium and self-similar over thirteen decades in time. Nat. Phys. 2016, 12, 171–174.

(77) Doxastakis, M.; Theodorou, D. N.; Fytas, G.; Kremer, F.; Faller, R.; Müller-Plathe, F.; Hadjichristidis, N. Chain and local dynamics of polyisoprene as probed by experiments and computer simulations. J. Chem. Phys. 2003, 119, 6883–6894.

(78) Szatkowski, L.; Merz, D. E.; Jiang, N.; Ejikeme, I.; Belonogov, L.; Ross, J. L.; Dima, R. I. Mechanics of the Microtubule Seam Interface Probed by Molecular Simulations and in Vitro Severing Experiments. J. Phys. Chem. B 2019, 123, 4888–4900.

(79) Varikoti, R. A.; Macke, A. C.; Speck, V.; Ross, J. L.; Dima, R. I. Molecular investigations into the unfoldase action of severing enzymes on microtubules. Cytoskeleton 2020, 77, 214–228.

(80) Lee, S.; Sowa, M. E.; hei Watanabe, Y.; Sigler, P. B.; Chiu, W.; Yoshida, M.; Tsai, F. T. The Structure of ClpB: A Molecular Chaperone that Rescues Proteins from an Aggregated State. Cell 2003, 115, 229–240.

(81) Luo, G.; Andricioaei, I.; Xie, X. S.; Karplus, M. Dynamic Distance Disorder in Proteins Is Caused by Trapping. J. Phys. Chem. B 2006, 110, 9363–9367, PMID: 16686476.

(82) Baysal, C.; Atilgan, A. R. Relaxation Kinetics and the Glassiness of Native Proteins: Coupling of Timescales. Biophys. J. 2005, 88, 1570–1576.

(83) Yang, H.; Luo, G.; Karnchanaphanurach, P.; Louie, T.-M.; Rech, I.; Cova, S.; Xun, L.; Xie, X. S. Protein Conformational Dynamics Probed by Single-Molecule Electron Transfer. Science 2003, 302, 262–266.

(84) Min, W.; Luo, G.; Cherayil, B. J.; Kou, S. C.; Xie, X. S. Observation of a Power-Law Memory Kernel for Fluctuations within a Single Protein Molecule. Phys. Rev. Lett. 2005, 94, 198302.

(85) Grossman-Haham, I.; Rosenblum, G.; Namani, T.; Hofmann, H. Slow domain reconfiguration causes power-law kinetics in a two-state enzyme. Proc. Natl. Acad. Sci. USA. 2018, 115, 513–518.

(86) Musafia, B.; Buchner, V.; Arad, D. Complex Salt Bridges in Proteins: Statistical Analysis of Structure and Function. J. Mol. Biol. 1995, 254, 761–770.

