## Supplemental Material for "Factors Underlying Asymmetric Dynamics of Disaggregase and Microtubule Severing AAA+ Machines"

#### Methods

##### Initial configuration

In the spiral conformation of severing proteins all six protomers (A to F) form a right-handed spiral with a ~ 5-6 Å translation and a ~ 60˚ twist for each protomer resulting in a ~ 40 Å gap between protomers A and F. Whereas, in the ring conformation protomer A is loosely bound to protomer B and interacts with protomer F, thereby closing the AAA+ ring. The substrate binds in a ~ 20 Å pore in the center of these hexameric conformations. The katanin hexamers, 6UGD (spiral) and 6UGE (ring), have two regions with missing residues: 183-187, and 324-331. The spastin hexamers, 6P07 (spiral) and 6PEN (ring), have the following C-terminal residues missing: 502-516, 613-619 and 752-758 in 6P07 and 415-420, 449-458, 473-477, 611-616 in 6PEN [Sandate C, 2019]. ATP is present in all 6 protomers from 6UGD and 6P07, while being absent in protomer A from 6UGE. 6PEN is the only structure solved in the presence of ADP in all the protomers but protomer F. Consequently, we used the Modeller program (version 9.23)

[Eswar N, 2008] to construct the missing residues in the above mentioned structures. We prepared 10 models for all the proteins using the standard parameters and ranked them based on their DOPE score. We chose one model for each protein based on the internal scoring function from Modeller and we used it as initial configuration-for our molecular dynamics simulations. Initial conformations of ClpB both lack N-terminal domains (amino acids 1-160) and middle domains (amino acids 409-524) of each protomer. Missing loop residues of the ring conformer (amino acid residues 525-528 in chain E and 284-293 in chain F) and of the spiral conformer (amino acid residues 284-294 in chains A-F and 684-659 in chain F) were modeled using the Modeller software using the approach described for severing proteins. A linker comprising five Gly residues was included to connect residues 408 and 525 in the absence of the middle domains. In the ring configuration (6OAX), ATP (ATP$\gamma$S) is present in all protomers except for ADP-bound NBD1 of protomer F and NBD2 of protomers A, F. In the spiral configuration (6OAY), ATP is bound to NBD2 of protomers C, D, E and NBD1 of protomer B and ADP is bound to NBD1 of protomers A and F and NBD2 of protomers A and B. NBD2 of protomer F is nucleotide free (APO) in the spiral conformer of ClpB.

#### Molecular Dynamics Simulations

The automated topology builder server was used to generate force field parameters for ATP, and ADP based on the GROMOS96 54A7 force field [Malde A, 2011]. The ATPase was placed at the center of a cubic box, with dimensions ~160x160x160 Å$^3$ for severing proteins and ~170x170x170 Å$^3$ for ClpB, solvated with water molecules represented using the single point charge (SPC) model, and neutralized by adding NaCl ions. Periodic boundary conditions (PBC) are applied in the three-dimensions. The number of particles of each system is shown in Table S3. To remove steric clashes, energy minimization was performed using the steepest descent algorithm and the Verlet cutoff-scheme for 50,000 steps with the convergence criterion of the maximum force value smaller than 23.9006 kcal/mol/A (1000 kJ/mol/nm). Next, MD simulations were performed to equilibrate the system and to obtain production data at T = 300 K. In the first equilibration step, to heat the system up to 300 K, simulations were performed in the NVT ensemble for 1 ns for severing proteins (500 ps for ClpB) using the leapfrog integrator and restraining the heavy atoms of the solute with harmonic forces with spring constant of 10 kcal·mol−1 Å−2. The second equilibration step of the constrained system was performed in the NPT ensemble for 1ns for severing proteins (500 ps for ClpB) by keeping pressure constant at 1.0 bar, using the Parrinello-Rahman pressure coupling scheme [Parrinello M, 1981], and temperature at 300 K, using the modified Berendsen thermostat with the response time of 0.1 ps [Bussi G, 2007]. Finally, five 50 ns production MD simulations were carried out without constraints. The bond lengths involving hydrogen atoms were restrained by using the LINCS method so that a longer integration step of 2 fs could be adopted. A distance cutoff of 10.0 Å was applied to compute the nonbonded interactions and-electrostatic interactions were calculated using the Particle Mesh Ewald (PME) algorithm [Daren T, 1993].

### Data Analysis

#### RMSF

To determine the contribution of each amino acid to the motion of our proteins, we calculated the root mean square fluctuation (RMSF) for each MD trajectory and we averaged over all the results from a given type of run. The RMSF calculation gives information about an individual residue flexibility by averaging the residue position over time (**Fig. 1**). The RMSF is defined as:

$$RMSF_i = \sqrt{\frac{1}{T}\sum_{t_j=1}^{N} |r_i(t_j) - r_i^{ref}|^2} \quad \text{.......................................(S1)}$$

$$< RMSF_i > \; = \frac{1}{n_{traj}} \sum_{k=1}^{n_{traj}} (RMSF_i)_k \quad \text{.....................................(S2)}$$

where N is the total number of frames of each trajectory, $r_i(t_j)$ is the position of the C-alpha atom of ith residue of the complex at time $t_j$ and $r_i^{ref}$ is the reference position of the i-th residue, $n_{traj}$ is the total number of trajectories and $<RMSF_i>$ is the average RMSF.

### Tables

**Table S1.** Details of the PDB structures for severing proteins used in our simulations.

| Protein | PDB ID | Source | Resolution (Å) | State | Substrate | Ligand |
|---|---|---|---|---|---|---|
| KATANIn | 6UGD | *Caenorhabditis elegans* | 3.5 | Spiral | Poly-Glutamate (E14) | ATP |
| | 6UGE | *Caenorhabditis elegans* | 3.6 | Ring | Poly-Glutamate (E12) | ATP |
| SPASTIN | 6P07 | *Drosophila melanogaster* | 3.2 | Spiral | Poly-Glutamate (E15) | ATP |
| | 6PEN | *Homo sapiens* | 4.2 | Ring | $(EY)_5$ | ADP |

**Table S2.** Details of the PDB structures for ClpB used in our simulations.

| Protein | PDB ID | Source | Resolution (Å) | State | Substrate | Ligand |
|---|---|---|---|---|---|---|
| ClpB | 6OAY | *Escherichia coli* | 3.3 | Spiral | Poly-Alanine (A26) | AGS/ADP |
| | 6OAX | *Escherichia coli* | 2.9 | Ring | Poly-Alanine (A26) | AGS/ADP |

**Table S3.** Summary of the MD setups for the 2 severing proteins in the various states.

| Protein | Nucleotide | Substrate | #Atoms | #Residues | #Water | #Na Ions | #Trajectories | Simulation time (ns) |
|---|---|---|---|---|---|---|---|---|
| **6UGD** | - | - | 419184 | 1902 | 400278 | 42 | 5 | 250 |
| | ATP | - | 419109 | 1902 | 399921 | 60 | 5 | 250 |
| | - | E14 | 419104 | 1916 | 400041 | 56 | 5 | 250 |
| | ATP | E14 | 419068 | 1916 | 399723 | 74 | 5 | 250 |
| **6UGE** | - | - | 420972 | 1902 | 402066 | 42 | 5 | 250 |
| | ATP | - | 420973 | 1902 | 401832 | 57 | 5 | 250 |
| | - | E12 | 420903 | 1914 | 401862 | 54 | 5 | 250 |
| | ATP | E12 | 420886 | 1914 | 401610 | 69 | 5 | 250 |
| **6P07** | - | - | 348393 | 1824 | 330471 | 24 | 5 | 250 |
| | ATP | - | 362364 | 1824 | 344160 | 42 | 5 | 250 |
| | - | E15 | 362349 | 1839 | 344259 | 39 | 5 | 250 |
| | ATP | E15 | 896625 | 1839 | 878253 | 57 | 5 | 250 |
| **6PEN** | - | - | 325244 | 1733 | 308139 | 0 | 5 | 250 |
| | ADP | - | 325190 | 1733 | 307875 | 15 | 5 | 250 |
| | - | (EY)5 | 325230 | 1743 | 307977 | 5 | 5 | 250 |
| | ADP | (EY)5 | 325134 | 1743 | 307671 | 20 | 5 | 250 |

**Table S4.** The summary of the details of the MD setups for the ClpB in the various states.

| Protein | Nucleotide | Substrate | #Atoms | #Residues | #Water | #Na Ions | #Trajectories | Simulation time (ns) |
|---|---|---|---|---|---|---|---|---|
| **6OAX** | - | - | 452280 | 3516 | 139088 | 72 | 5 | 250 |
| | AGS/ADP | - | 452304 | 3516 | 138913 | 105 | 5 | 250 |
| | - | poly-alanine | 452283 | 3516 | 139036 | 72 | 5 | 250 |
| | AGS/ADP | poly-alanine | 452253 | 3516 | 138948 | 105 | 5 | 250 |
| **6OAY** | - | - | 452019 | 3516 | 139001 | 72 | 5 | 250 |
| | AGS/ADP | - | 485282 | 3516 | 149923 | 101 | 5 | 250 |
| | - | poly-alanine | 452023 | 3516 | 138956 | 72 | 5 | 250 |
| | AGS/ADP | poly-alanine | 452015 | 3516 | 138781 | 101 | 5 | 250 |

**Table S5:** The list of the specific inter- and intra-protomer salt-bridges present in more than 3 protomers for longer than 1 ns in the various setups from the MD simulations of the severing proteins.

| **Present in >3 protomers [ > 1ns]** | | |
|---|---|---|
| **Setup** | **Inter** | **Intra** |
| **6UGD APO** | D171-K265 [BA, CB, DC, ED, FE] | R267-E308 [AA, BB, DD, EE, FF] |
| | D269-K265 [BA, CB, DC, ED, FE] | R301-D346 [AA, BB, CC, DD, EE, FF] |
| | R275-D261 [BA, CB, DC, ED, FE] | K314-D346 [AA, BB, CC, EE, FF] |
| | R301-E293 [BA, CB, DC, FE] | |
| | R311-E344 [CB, DC, ED, FE] | |
| **E14** | D171-K265 [BA, CB, DC, ED] | D269-K272 [AA, CC, EE, FF] |
| | D269-K265 [BA, CB, DC, ED, FE] | R301-D346 [AA, BB, CC, DD, EE, FF] |
| | R275-D261 [BA, CB, DC, ED, FE] | K314-D346 [BB, CC, DD, EE, FF] |
| | R301-E293 [BA, CB, DC, ED, FE] | |
| | R311-E344 [DC, ED, FE] | |
| **ATP** | D171-K265 [BA, DC, ED, FE] | R267-E308 [BB, CC, DD, EE, FF] |
| | D269-K265 [BA, CB, DC, ED, FE] | R301-D346 [AA, BB, CC, DD, EE, FF] |
| | R275-D261 [BA, CB, DC, ED, FE] | K314-D346 [AA, BB, EE, FF] |
| | R301-E293 [BA, CB, DC, ED, FE] | |
| | R311-E344 [CB, DC, ED, FE] | |
| **ATP +E14** | D171-K265 [BA, DC, FE] | D269-K272 [AA, BB, CC, DD, EE, FF] |
| | D269-K265 [BA, CB, DC, ED, FE] | R267-E308 [AA, BB, CC, DD] |
| | R275-D261 [BA, CB, DC, ED, FE] | R301-D346 [AA, BB, CC, DD, EE, FF] |
| | R301-E293 [BA, CB, DC, ED, FE] | K314-D346 [AA, BB, CC, DD, EE, FF] |

| **Present in >3 protomers [ > 1 ns]** | | |
|---|---|---|
| **Setup** | **Inter** | **Intra** |
| **6UGE APO** | D171-K265 [DC, ED, FE] | D269-K272 [AA, BB, DD] |
| | D269-K265 [BA, CB, DC, ED, FE] | R301-D346 [AA, BB, CC, DD, EE, FF] |
| | R275-D261 [CB, DC, ED, FE] | K314-D346 [AA, BB, CC, DD, EE, FF] |
| | R301-E293 [CB, ED, FE] | |
| | R301-D295 [DC, ED, FE] | |

| | | |
|---|---|---|
| | R301-E344 [BA, CB, ED, FE] | |
| **E12** | D171-K265 [CB, ED, FE] | D269-K272 [AA, BB, CC, DD] |
| | D269-K265 [CB, DC, ED, FE] | R267-E308 [AA, BB, CC, DD, EE] |
| | R275-D261 [CB, DC, ED, FE] | R301-D346 [AA, BB, CC, DD, EE] |
| | R301-E293 [CB, DC, ED, FE] | K314-D346 [AA, BB, CC, DD, EE, FF] |
| **ATP** | D171-K265 [CB, DC, ED, FE] | D269-K272 [AA, BB, CC, EE, FF] |
| | D269-K265 [CB, DC, ED, FE] | R267-E308 [AA, CC, DD, EE, FF] |
| | R275-D261 [CB, DC, ED, FE] | R301-D346 [AA, BB, CC, DD, EE, FF] |
| | R301-E293 [DC, ED, FE] | K314-D346 [AA, BB, CC, DD, EE, FF] |
| | R301-D295 [CB, ED, FE] | |
| | R301-E344 [BA, CB, DC, FE] | |
| **ATP + E12** | D269-K265 [CB, DC, ED, FE] | D269-K272 [AA, BB, CC, EE] |
| | R275-D261 [CB, DC, ED, FE] | R301-D346 [AA, BB, CC, DD, EE, FF] |
| | | K314-D346 [AA, BB, CC, DD, EE, FF] |

| **Present in >3 protomers [ > 1 ns]** | | |
|---|---|---|
| **Setup** | **Inter** | **Intra** |
| **6P07 APO** | E462-K555 [BA, CB, DC, ED, FE] | E561-R601 [BB, CC, EE, FF] |
| | D559-K555 [BA, CB, DC, ED, FE] | R591-D635 [BB, CC, DD, EE, FF] |
| | R591-D585 [BA, DC, ED, FE] | E595-K603 [AA, BB, CC, DD, EE, FF] |
| | R591-E633 [BA, CB, DC, FE] | |
| **E15** | D559-K555[BA, CB, DC, ED, FE] | E561-R601 [AA, BB, CC, EE, FF] |
| | R591-D585 [BA, CB, ED, FE] | R591-D635 [AA, BB, CC, DD, EE, FF] |
| | | E595-K603 [AA, BB, CC, DD, EE, FF] |
| **ATP** | D559-K555 [BA, CB, DC, ED, FE] | R591-D635 [AA, BB, CC, DD, EE, FF] |
| | R591-D585 [BA, CB, ED, FE] | E595-K603 [AA, BB, CC, DD, EE, FF] |
| | | K603-D635 [AA, BB, CC, DD, EE, FF] |
| **ATP + E15** | E462-K555 [DC, ED, FE] | E561-R601 [AA, DD, EE] |
| | D559-K555 [BA, CB, DC, ED, FE] | R591-D635 [AA, BB, CC, DD, EE, FF] |
| | R591-D585 [BA, CB, DC, ED, FE] | E595-K603 [AA, BB, CC, DD, EE, FF] |
| | R591-E633 [CB, DC, FE] | K603-D635 [AA, BB, CC, DD, EE] |
| | R600-E633 [BA, CB, FE] | |

| **Present in >3 protomers [ > 1 ns]** | | |
|---|---|---|
| **Setup** | **Inter** | **Intra** |
| **6PEN APO** | E418-K414 [BA, CB, DC, ED] | E420-R460 [AA, BB, CC, DD, EE] |
| | K421-E331 [BA, CB, DC, ED] | R450-D493 [AA, BB, CC, DD, EE] |
| | R450-D444 [BA, CB, DC] | E454-R459 [AA, BB, CC, DD, EE] |
| | R450-E491 [BA, CB, ED] | E454-K462 [AA, BB, CC, DD, EE] |
| | | D456-R460 [AA, BB, CC, DD, EE] |
| | | K462-D493 [AA, BB, CC, DD, EE, FF] |
| | | E494-R498 [AA, BB, CC, DD, EE, FF] |
| **EY5** | E418-K414 [BA, CB, DC, ED] | E420-R460 [AA, BB, CC, DD, EE] |
| | R450-D444 [BA, CB, DC, ED, AF] | R450-D493 [AA, BB, CC, DD, EE] |
| | D456-K462 [BA, CB, DC, ED] | E454-R459 [AA, BB, CC, DD] |
| | | E454-K462 [AA, BB, CC, DD, EE] |
| | | D456-R460 [AA, BB, CC, EE, FF] |
| | | K462-D493 [AA, BB, CC, DD, EE, FF] |
| | | E494-R498 [AA, BB, CC, DD, EE, FF] |
| **ADP** | E418-K414 [BA, CB, DC, ED] | E420-R460 [AA, BB, CC, DD, EE] |
| | K421-E331 [BA, CB, DC, ED] | D493-R460 [BB, CC, DD, EE] |
| | R450-E442 [BA, CB, DC, ED, AF] | E454-R459 [BB, DD, EE] |
| | R450-E491 [BA, CB, ED] | E454-K462 [AA, BB, CC, DD, EE] |
| | D456-K462 [CB, DC, ED] | D456-R460 [AA, BB, CC, DD, EE] |
| | | K462-D493 [BB, CC, DD, EE, FF] |
| | | E494-R498 [AA, BB, DD, EE, FF] |
| **ADP + EY5** | E418-K414 [BA, CB, DC, ED] | E420-R460 [AA, BB, CC] |
| | R450-E442 [BA, CB, DC, ED, AF] | R450-D493 [AA, BB, CC, DD, EE] |
| | D456-K462 [BA, CB, DC, ED] | R451-E491 [CC, DD, EE] |
| | | E454-K462 [AA, BB, CC, DD, EE] |

| | | |
|---|---|---|
| | | D456-R460 [AA, BB, CC, DD, EE, FF] |
| | | K462-D493 [AA, CC, DD, EE, FF] |
| | | E494-R498 [AA, BB, EE, FF] |

**Table S6:** The list of the specific inter- and intra-protomer salt-bridges present in more than 3 protomers for longer than 1 ns in the various setups from the MD simulations of ClpB

| **Present in >3 protomers [ > 1 ns]** | | |
|---|---|---|
| **Setup** | **Inter** | **Intra** |
| **6OAX APO** | E639-K640 [BA, CB, ED] | E636-R645[AA, BB, CC, DD, EE]<br>D290-R252[CC, EE, FF] |
| | D290-K288[AB, BC, DE, EF] | |
| | E254-K250[AB, BC, DE] | |
| | E257-K250[AB, BC, DE] | |
| **ALA** | D290-K288[AB, BC, CD, DE, FE] | E636-R645[BB. CC, EE] |
| | E254-K250[AB, BC, DE] | |
| | E256-R252[BA, CB, DC] | |
| | E639-K640[AF, BA, CB, ED] | |
| | E257-K250[AB, BC, DE] | |
| **ATP** | D290-K288[AB, DE] | E636-R645[BB, CC, DD, EE] |
| | E639-K640[CB, DC, ED, FE] | |
| | E254-K250[AB, BC, DE] | |
| **ALA+ATP** | E254-K250[AB, BC, DE] | E636-R645[BB, CC, DD, EE]<br>D290-R252[CC, EE, FF] |
| | E639-K640[AF, BA, CB, DC, ED, FE] | |
| | E257-K250[AB, BC, DE] | |

| **Present in >3 protomers [ > 1 ns]** | | |
|---|---|---|
| **Setup** | **Inter** | **Intra** |
| **6OAY APO** | E636-R645[AF, BA, DC, ED] | E254-R258[AA, BB, EE] |
| | E254-K250[AB, BC, CD, EF, FA] | |
| | E639-K640[AF, BA, CB, DC, FE] | |
| | E257-K250[AB, BC, CD, DE, FA] | |

| | | |
|---|---|---|
| **ALA** | E254-K250[BC, CD, DC, DE, ED, FE] | E254-R258[BB, CC, EE, FF] |
| | E639-K640[CB, DC, ED] | E636-R645[BB, CC, DD, EE] |
| | E257-K250[BC, CD, DE] | |
| **ATP** | E257-K250[AB, BC, CD, DE] | E636-R645[AA, BB, DD, FF] |
| | E639-K640[AF, BA, CB, DC, ED] | |
| | D290-K250[DE, EF, FA] | |
| | E254-K250[AB, BC, CD] | |
| **ALA+ATP** | E254-K250[BC, CD, DE] | |
| | E639-K640[AF, BA, CB, ED, FA] | E636-K645[BB, CC, EE] |
| | D290-K250[AB, CB, EF, FA] | E256-R252[BB, CC, EE] |
| | E257-K250[BC, DE, EF] | E254-K250[CC, EE, FF] |

**Table S7:** Parameters obtained from the fitting of the ACF data from the 50 ns long simulation trajectories for severing proteins with **Eq. 3** from the main text. The characteristic relaxation time, $\tau^*$, is calculated using **Eq. 4** from the main text.

| PDB | ns | $\beta$ | T1 (ns) | T2 (ns) | $\alpha$ | $\tau^*$ (ns) |
|---|---|---|---|---|---|---|
| **6UGD APO** | 50 | 0.5 | 0.082 | 6.28 | 0.279 | 4.57 |
| **6UGD ATP** | 50 | 0.5 | 0.145 | 5.00 | 0.387 | 3.40 |
| **6UGD E14** | 50 | 0.4 | 0.703 | 5.67 | 0.477 | 4.14 |
| **6UGD ATP+E14** | 50 | 0.5 | 0.156 | 6.01 | 0.413 | 3.60 |
| **6UGE APO** | 50 | 0.4 | 0.089 | 4.93 | 0.332 | 3.50 |
| **6UGE ATP** | 50 | 0.5 | 0.083 | 4.95 | 0.365 | 3.27 |
| **6UGE E12** | 50 | 0.4 | 0.115 | 5.44 | 0.441 | 3.34 |
| **6UGE ATP+E12** | 50 | 0.4 | 0.050 | 4.30 | 0.338 | 3.11 |
| **6P07 APO** | 50 | 0.3 | 0.172 | 5.14 | 0.566 | 3.22 |
| **6P07 ATP** | 50 | 0.4 | 0.121 | 3.96 | 0.505 | 2.46 |
| **6P07 E15** | 50 | 0.5 | 1.577 | 3.70 | 0.582 | 3.85 |
| **6P07 ATP+E15** | 50 | 0.4 | 0.612 | 3.81 | 0.497 | 3.04 |
| **6PEN APO** | 50 | 0.4 | 0.208 | 6.90 | 0.434 | 4.67 |
| **6PEN ADP** | 50 | 0.5 | 1.106 | 3.56 | 0.498 | 2.74 |
| **6P07 (EY)5** | 50 | 0.4 | 0.064 | 5.05 | 0.472 | 2.97 |
| **6P07 ATP+(EY)5** | 50 | 0.4 | 0.076 | 3.51 | 0.472 | 2.74 |

**Table S8.** Parameters obtained from the fitting of the ACF data from the 5 ns long simulation trajectories for severing proteins with **Eq. 3** from the main text. The characteristic relaxation time, $\tau^*$, is calculated using **Eq. 4** from the main text.

| PDB | ns | $\beta$ | T1 (ns) | T2 (ns) | $\alpha$ | $\tau^*$ (ns) |
|---|---|---|---|---|---|---|
| **6UGD APO** | 5 | 0.5 | 0.023 | 0.69 | 0.352 | 0.48 |
| **6UGD ATP** | 5 | 0.5 | 0.029 | 0.53 | 0.361 | 0.39 |
| **6UGD E14** | 5 | 0.3 | 0.029 | 0.62 | 0.389 | 0.49 |
| **6UGD ATP+E14** | 5 | 0.5 | 0.012 | 0.52 | 0.330 | 0.39 |
| **6UGE APO** | 5 | 0.5 | 0.023 | 0.61 | 0.400 | 0.41 |
| **6UGE ATP** | 5 | 0.5 | 0.016 | 0.58 | 0.328 | 0.39 |
| **6UGE E12** | 5 | 0.5 | 0.010 | 0.51 | 0.386 | 0.37 |
| **6UGE ATP+E12** | 5 | 0.5 | 0.011 | 0.62 | 0.312 | 0.44 |
| **6P07 APO** | 5 | 0.4 | 0.011 | 0.48 | 0.432 | 0.29 |
| **6P07 ATP** | 5 | 0.5 | 0.047 | 0.70 | 0.428 | 0.44 |
| **6P07 E15** | 5 | 0.5 | 0.085 | 0.43 | 0.490 | 0.33 |
| **6P07 ATP+E15** | 5 | 0.5 | 0.167 | 0.29 | 0.535 | 0.35 |
| **6PEN APO** | 5 | 0.4 | 0.020 | 0.60 | 0.502 | 0.36 |
| **6PEN ADP** | 5 | 0.5 | 0.082 | 0.41 | 0.425 | 0.33 |
| **6P07 (EY)5** | 5 | 0.4 | 0.011 | 0.46 | 0.408 | 0.35 |
| **6P07 ATP+(EY)5** | 5 | 0.5 | 0.079 | 0.46 | 0.492 | 0.33 |

**Table S9:** Fitting parameters obtained from the stretched exponential plus single exponential fitting (**Eq. 3** in main text) performed on the ACF data derived from the 50 ns long simulation trajectories. The $\tau^*$ is calculated using **Eq. 4** (in main text).

| PDB | ns | $\beta$ | T1 (ns) | T2 (ns) | $\alpha$ | $\tau^*$ (ns) |
|---|---|---|---|---|---|---|
| **6OAX APO** | 50 | 0.5 | 0.052 | 4.42 | 0.200 | 3.39 |
| **6OAX ATP** | 50 | 0.5 | 0.595 | 4.45 | 0.400 | 4.21 |
| **6OAX ALA** | 50 | 0.6 | 1.836 | 4.55 | 0.372 | 4.03 |
| **6OAX ATP+ALA** | 50 | 0.4 | 0.114 | 5.33 | 0.400 | 3.57 |
| **6OAY APO** | 50 | 0.6 | 0.058 | 3.99 | 0.300 | 2.98 |
| **6OAY ATP** | 50 | 0.6 | 0.112 | 3.79 | 0.300 | 2.80 |
| **6OAY ALA** | 50 | 0.4 | 0.415 | 3.88 | 0.500 | 3.54 |
| **6OAY ATP+ALA** | 50 | 0.7 | 1.305 | 3.921 | 0.400 | 3.85 |

**Table S10.** Fitting parameters obtained from Stretched exponential plus single exponential fitting (**Eq. 3** in main text) performed on ACF data derived from short simulation times ($t$=5ns). The $\tau$* calculated using **Eq. 4** (in main text).

| PDB | ns | $\beta$ | T1 (ns) | T2 (ns) | $\alpha$ | $\tau$* (ns) |
|---|---|---|---|---|---|---|
| **6OAX APO** | 5 | 0.5 | 0.037 | 0.63 | 0.400 | 0.46 |
| **6OAX ATP** | 5 | 0.5 | 0.054 | 0.44 | 0.300 | 0.34 |
| **6OAX ALA** | 5 | 0.5 | 0.003 | 0.40 | 0.283 | 0.29 |
| **6OAX ATP+ALA** | 5 | 0.5 | 0.039 | 0.38 | 0.400 | 0.35 |
| **6OAY APO** | 5 | 0.5 | 0.069 | 0.39 | 0.300 | 0.43 |
| **6OAY ATP** | 5 | 0.6 | 0.007 | 0.48 | 0.200 | 0.40 |
| **6OAY ALA** | 5 | 0.5 | 0.014 | 0.43 | 0.300 | 0.28 |
| **6OAY ATP+ALA** | 5 | 0.7 | 0.164 | 0.33 | 0.300 | 0.37 |

**Table S11:** Extrapolation of $\tau$* for severing proteins to experimental timescales (1 millisecond) using the power law equation $\tau*(t) = b * t^{\theta}$ (shown in **Figure S12**).

| Setup | $\theta$ | b | t (millisecond) | $\tau$* (microseconds) |
|---|---|---|---|---|
| **6UGD APO** | 1.0815 | 0.066 | 1 | 203.49 |
| **6UGD ATP** | 1.00634 | 0.066 | 1 | 72.042 |
| **6UGD E14** | 1.12235 | 0.051 | 1 | 276.48 |
| **6UGD E14+ATP** | 1.07678 | 0.053 | 1 | 153.09 |
| **6UGE APO** | 1.10917 | 0.045 | 1 | 203.34 |
| **6UGE ATP** | 0.84641 | 0.12 | 1 | 14.38 |
| **6UGE E12** | 1.03594 | 0.058 | 1 | 95.29 |
| **6UGE E12+ATP** | 0.87862 | 0.1 | 1 | 18.69 |
| **6P07 APO** | 1.16021 | 0.034 | 1 | 310.98 |
| **6P07 ATP** | 0.71772 | 0.149 | 1 | 3.017 |
| **6P07 E15** | 1.10333 | 0.051 | 1 | 212.59 |
| **6P07 E15+ATP** | 0.88232 | 0.097 | 1 | 19.08 |
| **6PEN APO** | 1.1406 | 0.054 | 1 | 376.7 |
| **6PEN ADP** | 0.89701 | 0.082 | 1 | 19.76 |
| **6PEN EY5** | 0.82851 | 0.118 | 1 | 11.04 |
| **6PEN EY5+ADP** | 1.06398 | 0.042 | 1 | 101.66 |

**Table S12:** Extrapolation of $\tau^*$ to experimental timescales (1 millisecond). The $\tau^*$ values are selected for theoretical prediction of $\tau^*$ at a given time by performing linear fitting using the power law equation $\tau^*(t) = b * t^{\theta}$ (shown in **Figure S13**).

| Setup | $\boldsymbol{\theta}$ | b | t (millisecond) | $\boldsymbol{\tau}$* (microseconds) |
|---|---|---|---|---|
| **6OAX APO** | 0.874 | 0.111 | 1 | 19.47 |
| **6OAX ATP** | 0.996 | 0.086 | 1 | 81.38 |
| **6OAX ALA** | 0.992 | 0.084 | 1 | 75.21 |
| **6OAX ALA+ATP** | 1.042 | 0.060 | 1 | 107.19 |
| **6OAY APO** | 0.756 | 0.157 | 1 | 5.39 |
| **6OAY ATP** | 0.805 | 0.146 | 1 | 9.87 |
| **6OAY ALA** | 0.985 | 0.076 | 1 | 61.78 |
| **6OAY ALA+ATP** | 1.037 | 0.065 | 1 | 108.37 |

### Figures

**a)**

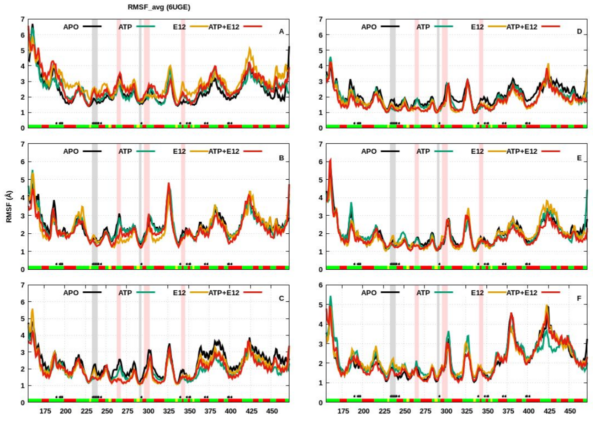


**b)**

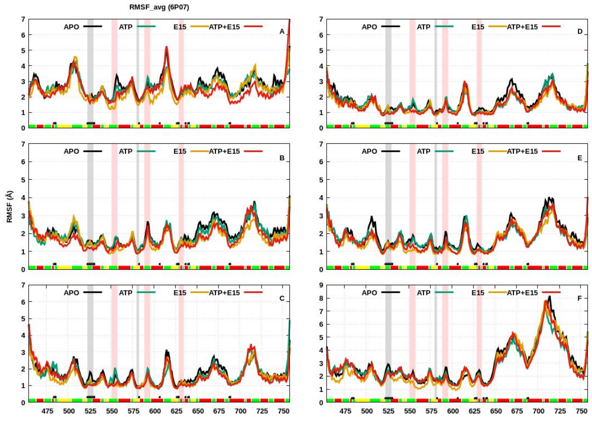


**c)**

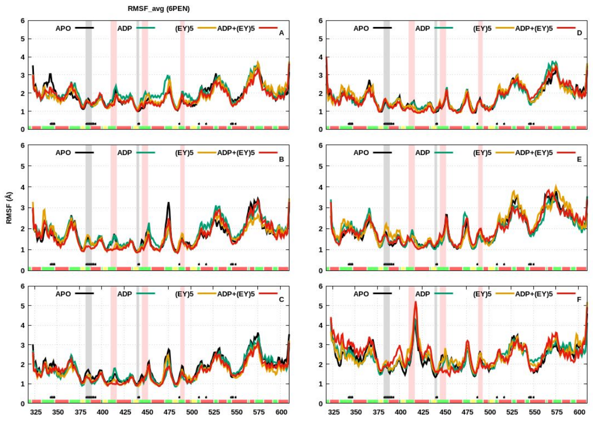


**Figure S1:** Average RMSF values for protomers in each state of the katanin and spastin severing proteins, as reported in **Fig. 1** from the main text. **a)** Ring katanin hexamer, **b)** Spiral spastin; in the APO state, ATP bound state, substrate bound state and substrate with ATP bound state, **c)** Ring spastin hexamer; in the APO state (black curve), ADP bound state (green), substrate bound state (orange) and substrate with ADP bound state (red). The secondary structures are shown below each figure above the x-axis. Different groups in the protein are annotated with shaded regions. Black dots (●) represent the ATP binding sites. Shaded regions in each plot show the functionally important elements: WA, WB (light gray), PL1, PL2 and PL3 (in light red).

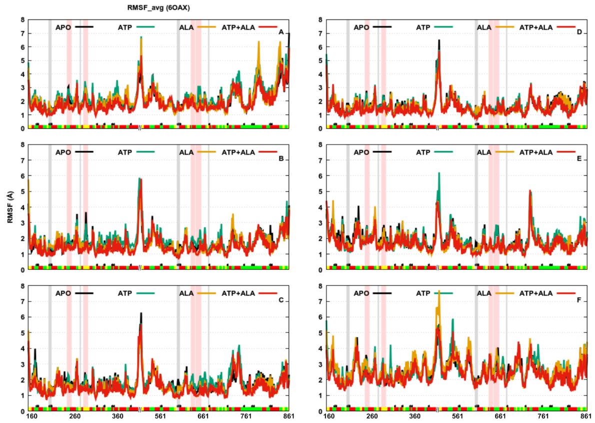


**Figure S2:** Average RMSF values for protomers in each state of ClpB, as reported in **Fig. 1** from the main text. Ring ClpB hexamer;in the APO state (black curve), ADP bound state (green), substrate bound state (orange) and substrate with ADP bound state (red). The secondary structures are shown below each figure above the x-axis. Different groups in the protein are annotated with shaded regions. Black dots (●) represent the ATP binding sites. Shaded regions in each plot show the functionally important elements: WA, WB (in light gray), PL1, PL2 and PL3 (in light red)

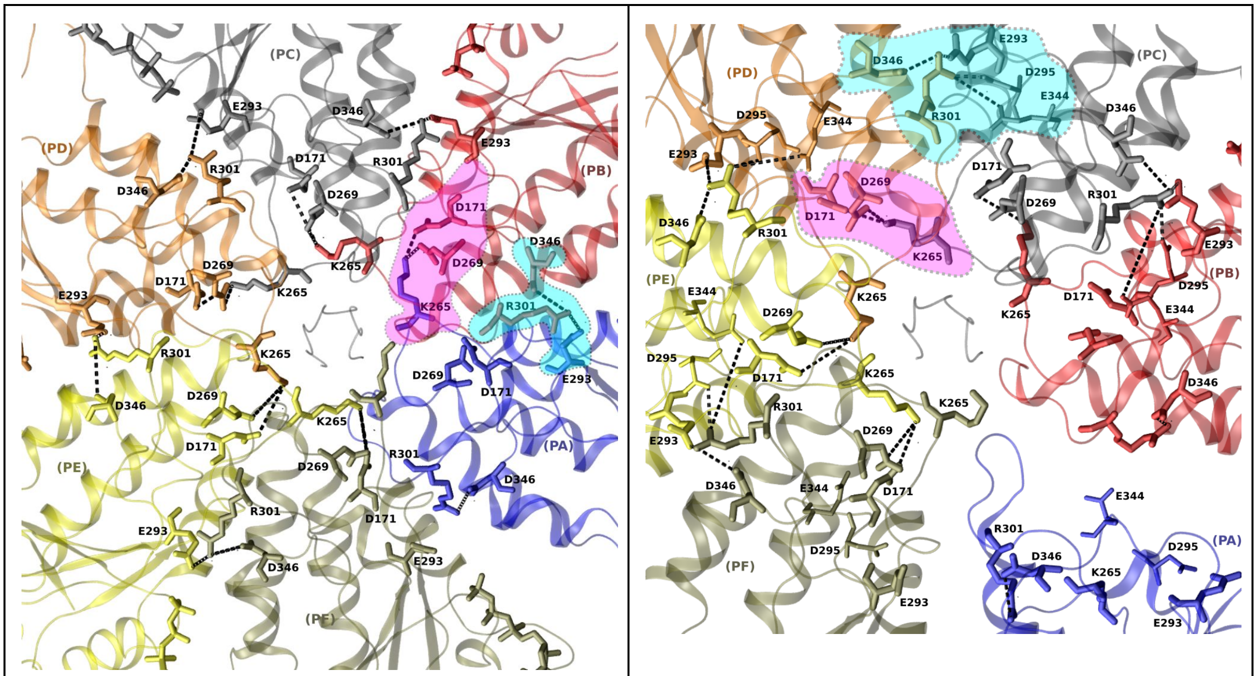


**Figure S3:** The two types of salt-bridge networks identified in our simulations for the katanin spiral (6UGD, left panel) and ring (6UGE, right panel): network (A) is in pink and network (B) in cyan. Each networks is shown for only one protomer pair.

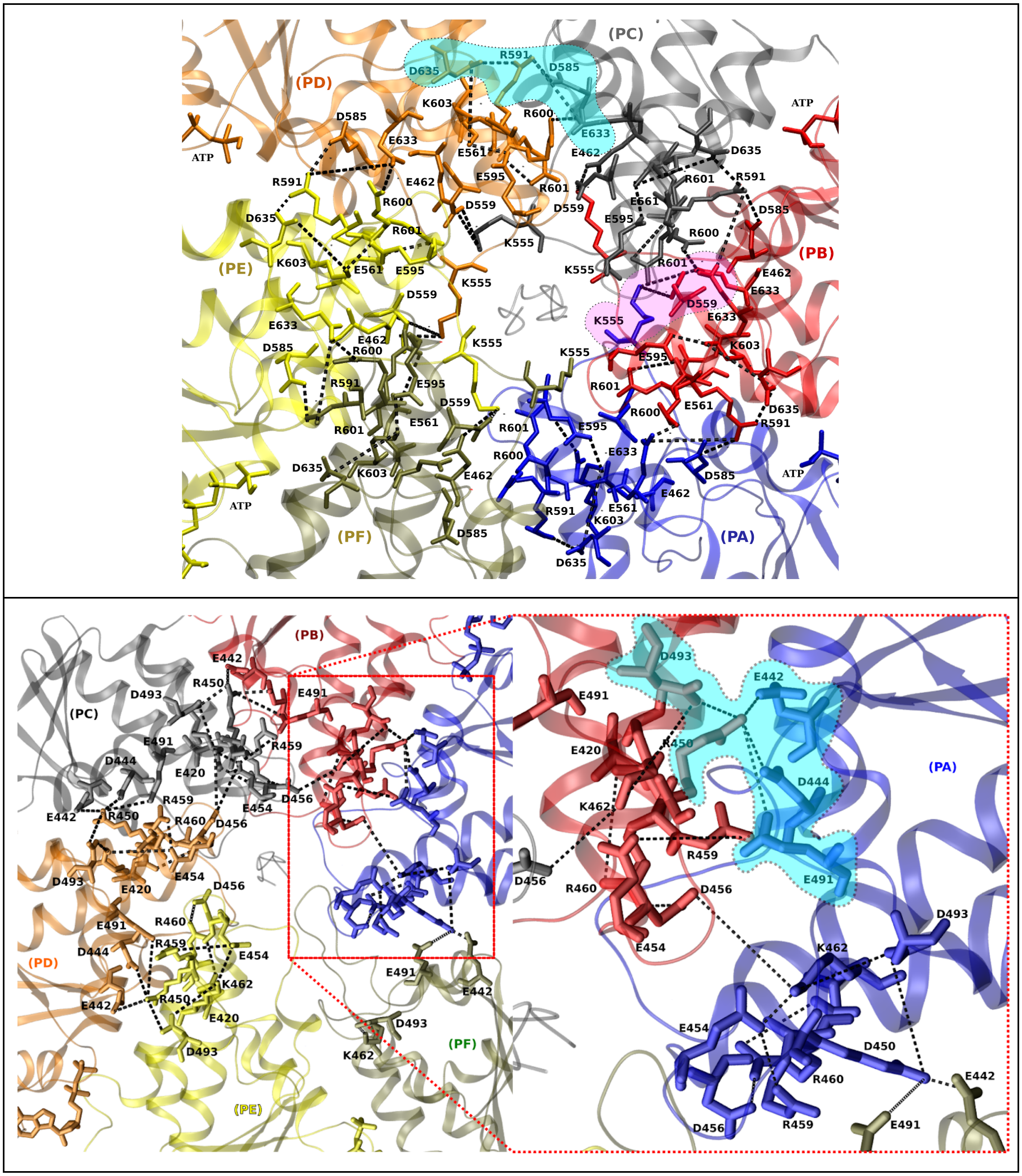

**Figure S4:** Similar to Figure S3 for the spastin spiral (6P07, upper panel) and ring (6PEN, lower panel).

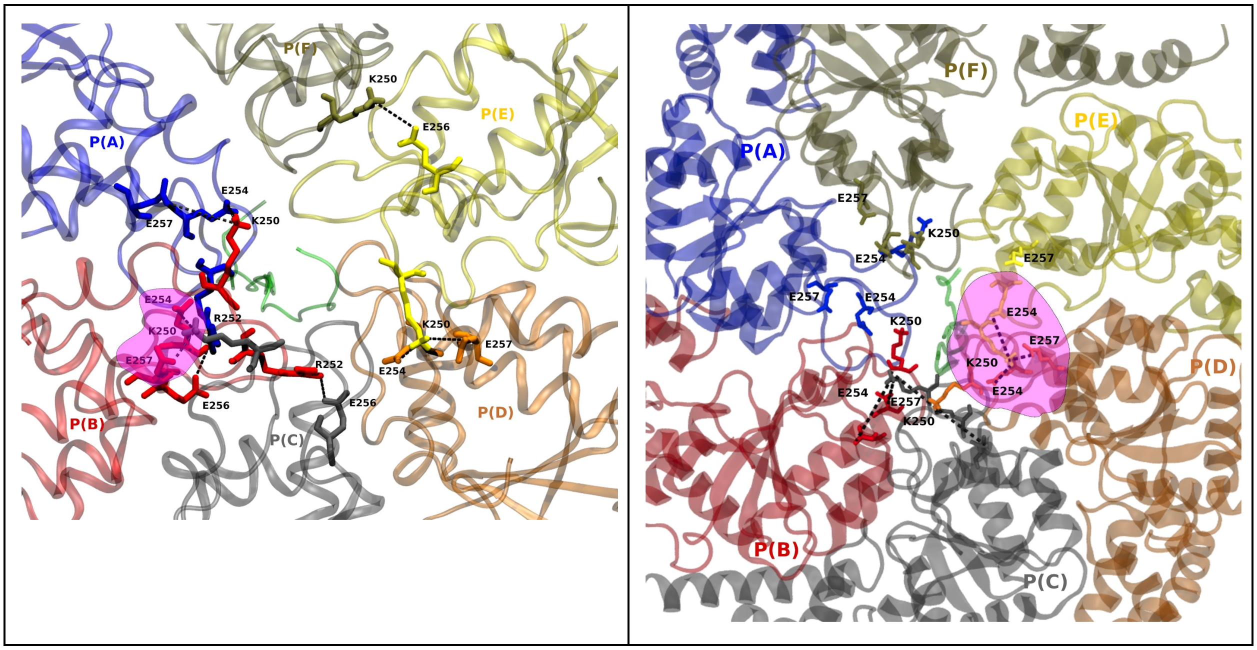


**Figure S5:** Types of salt-bridge networks identified in our simulations for the ClpB ring (6OAX, left panel), and spiral (6OAY, right panel): network formed by PL1 loops in pink (shown for only one protomer pair).

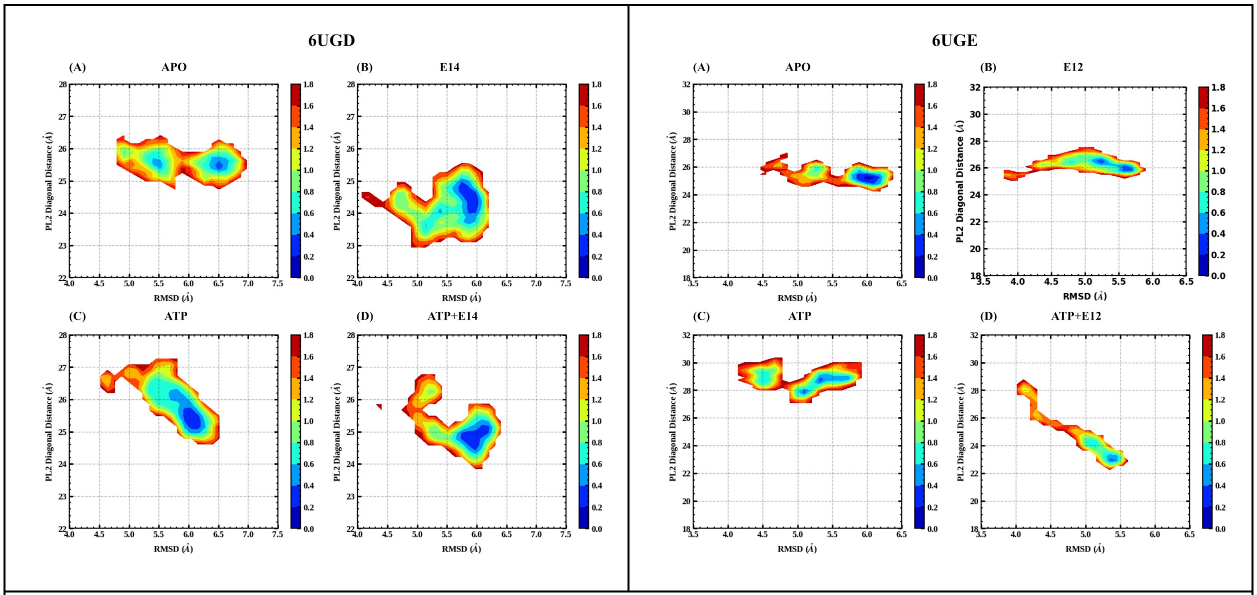


**Figure S6:** Free Energy Landscape plotted as a function of RMSD and PL2-diagonal distances in the katanin spiral (6UGD) and ring (6UGE).

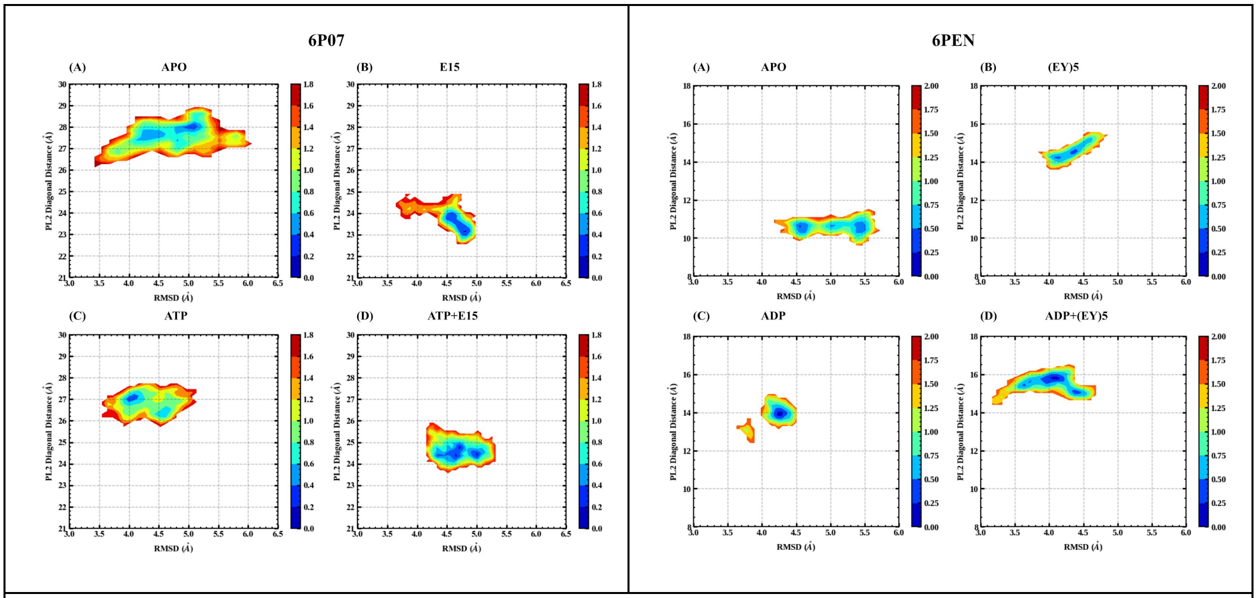


**Figure S7:** Similar to Figure S6 for the spastin spiral (6P07) and ring (6PEN).

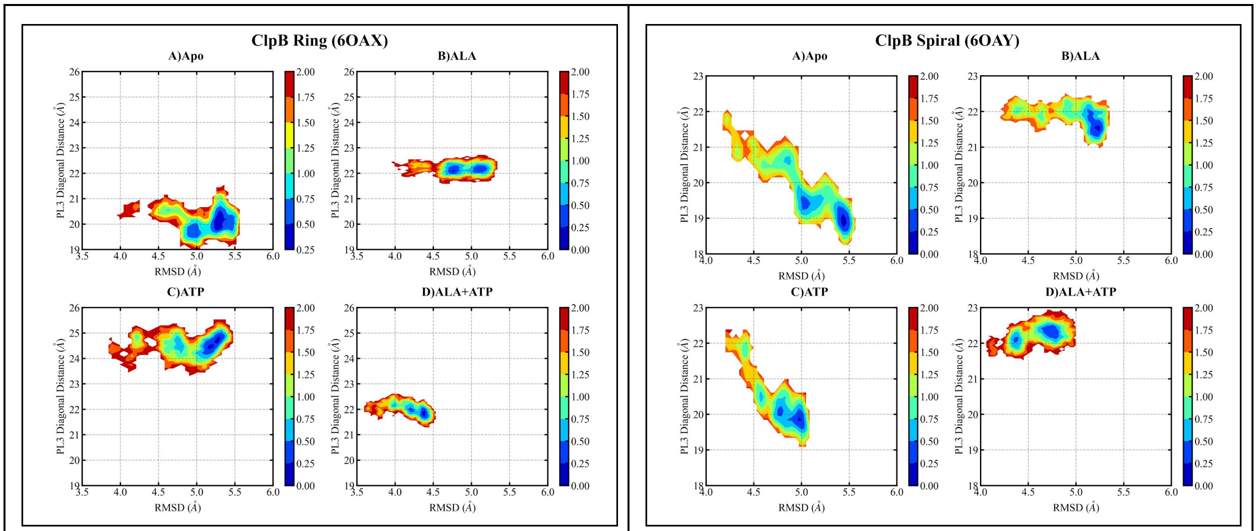


**Figure S8:** Free Energy Landscape plotted as a function of RMSD and PL3-diagonal distances in the ClpB ring (6OAX) and spiral (6OAY).

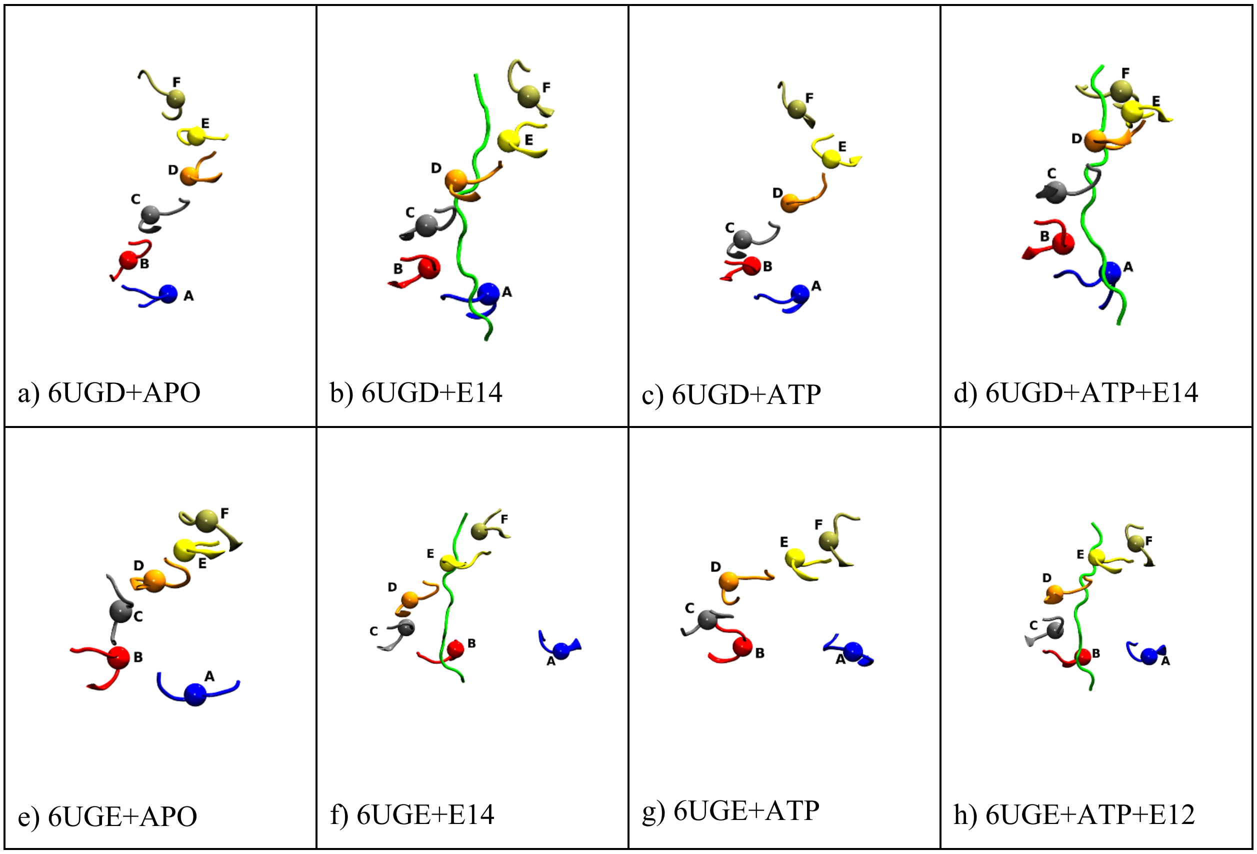


**Figure S9:** The PL1 loop arrangement in the katanin spiral (6UGD) and ring (6UGE) for the most populated state from each of the FEL plots from Figure **2** in main text. The loop from each protomer is colored differently: A blue, B red, C dark grey, D orange, E yellow, and F brown. The substrate peptide is indicated in green.

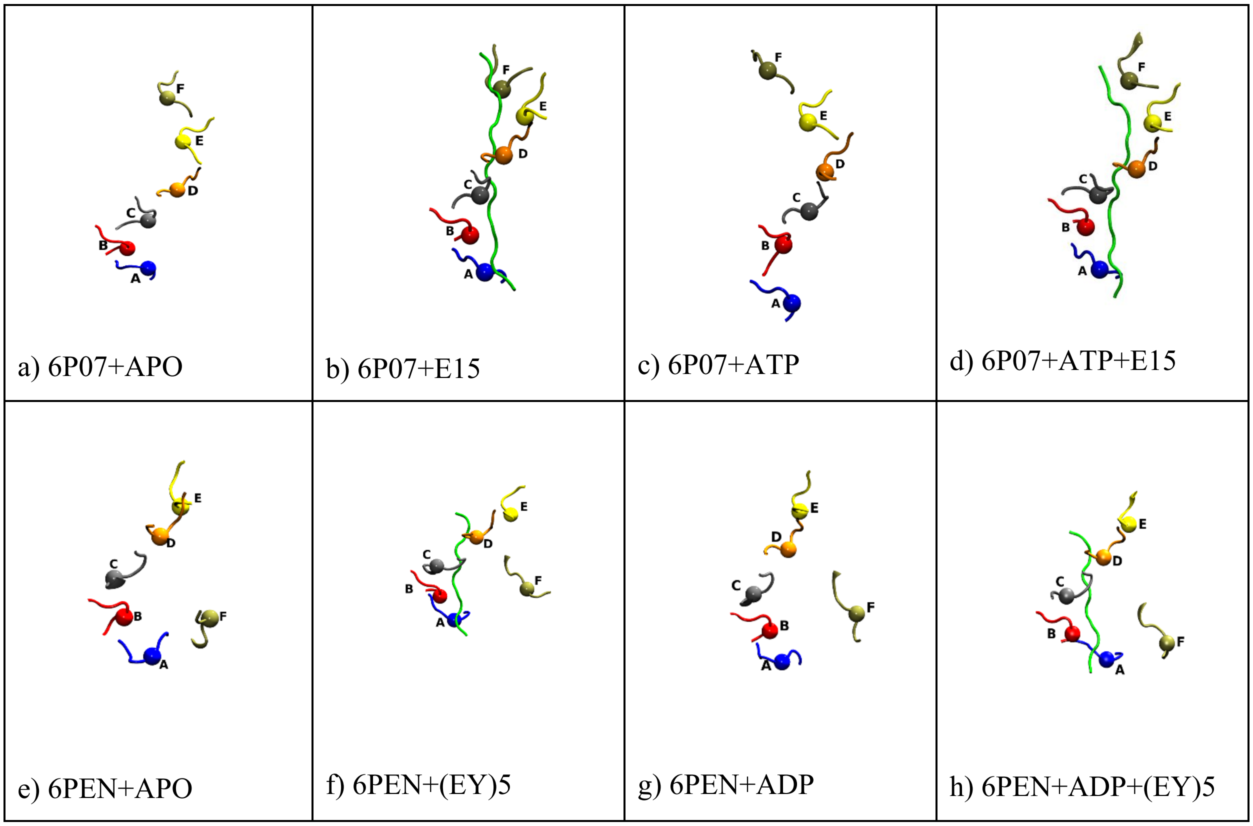


**Figure S10:** Similar to Figure S9 for spastin spiral (6P07) and ring (6PEN).

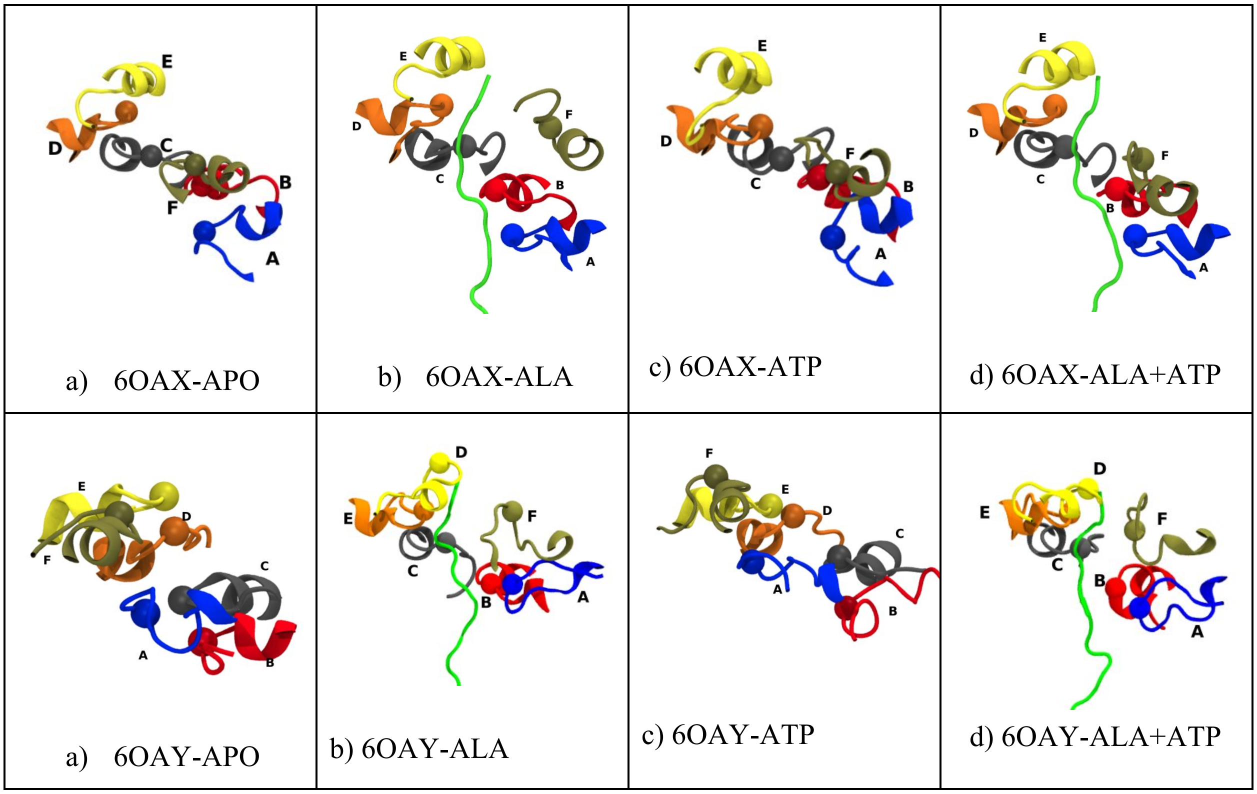


**Figure S11:** Similar to Figure S9 for ClpB ring (6OAX) and spiral (6OAY) for the most populated state from each of the FEL plots from Figure **3** in main text.

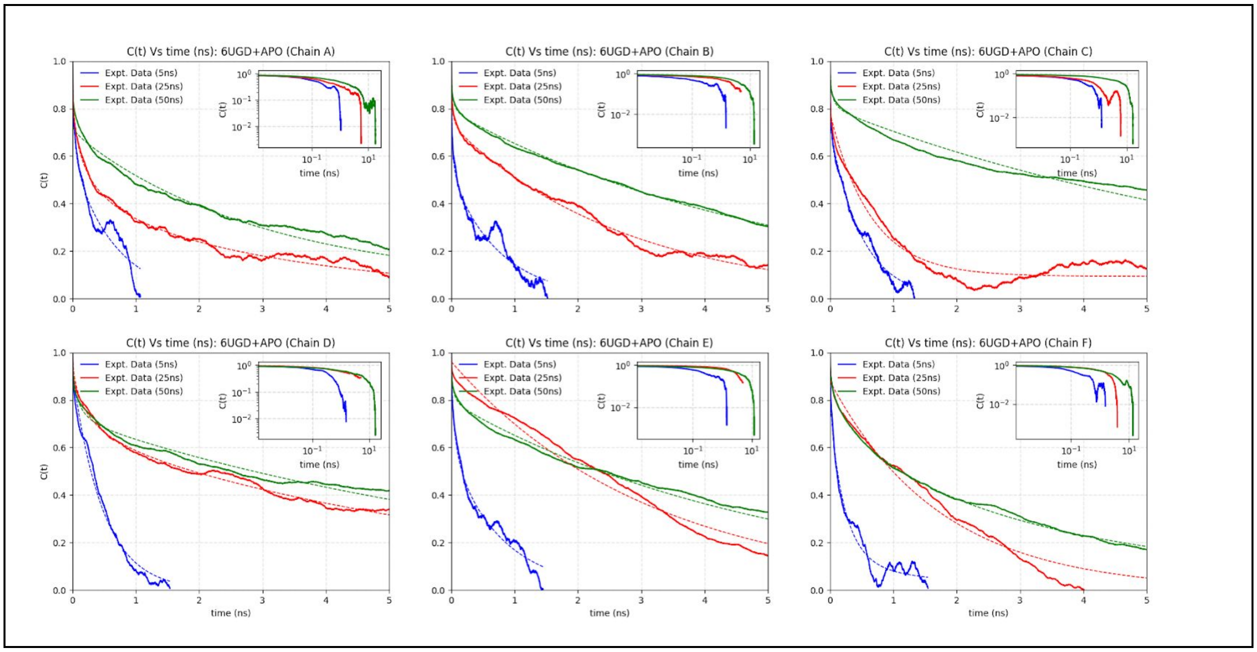


**Figure S12:** Sample relaxation curves for ACF in each protomer from the katanin spiral (6UGD) in the APO state: each panel shows C(t) obtained from 5 ns (blue), 25 ns (red) and 50 ns (green). The dashed lines represent fitting curves using **Eq. 3** from the main text. The same data are plotted on the log-log scale in the inset.

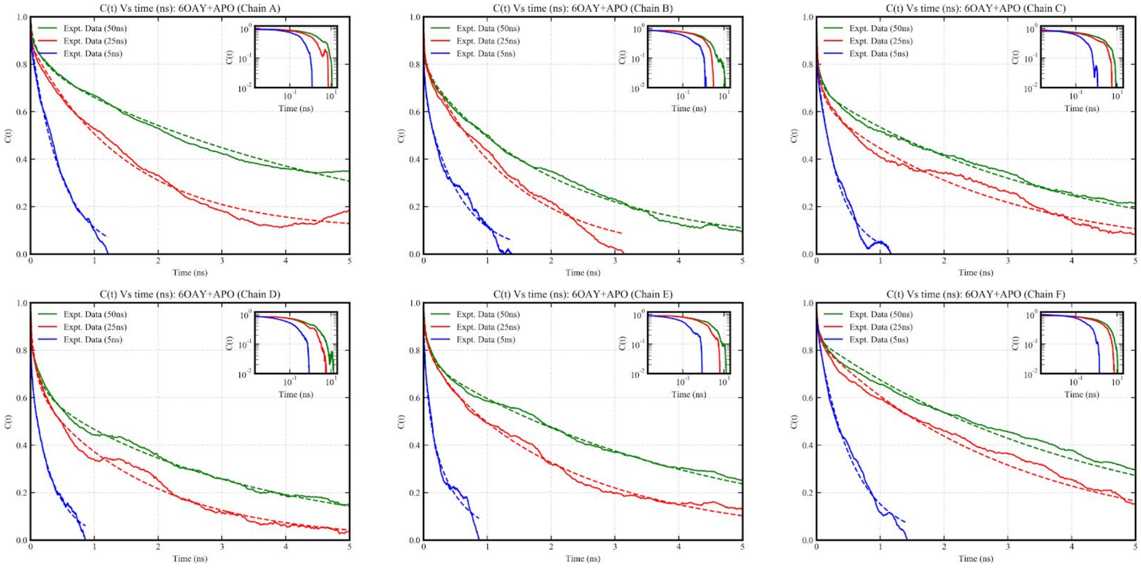


**Figure S13:** Similar to Figure S12 for ClpB spiral (6OAY) APO.

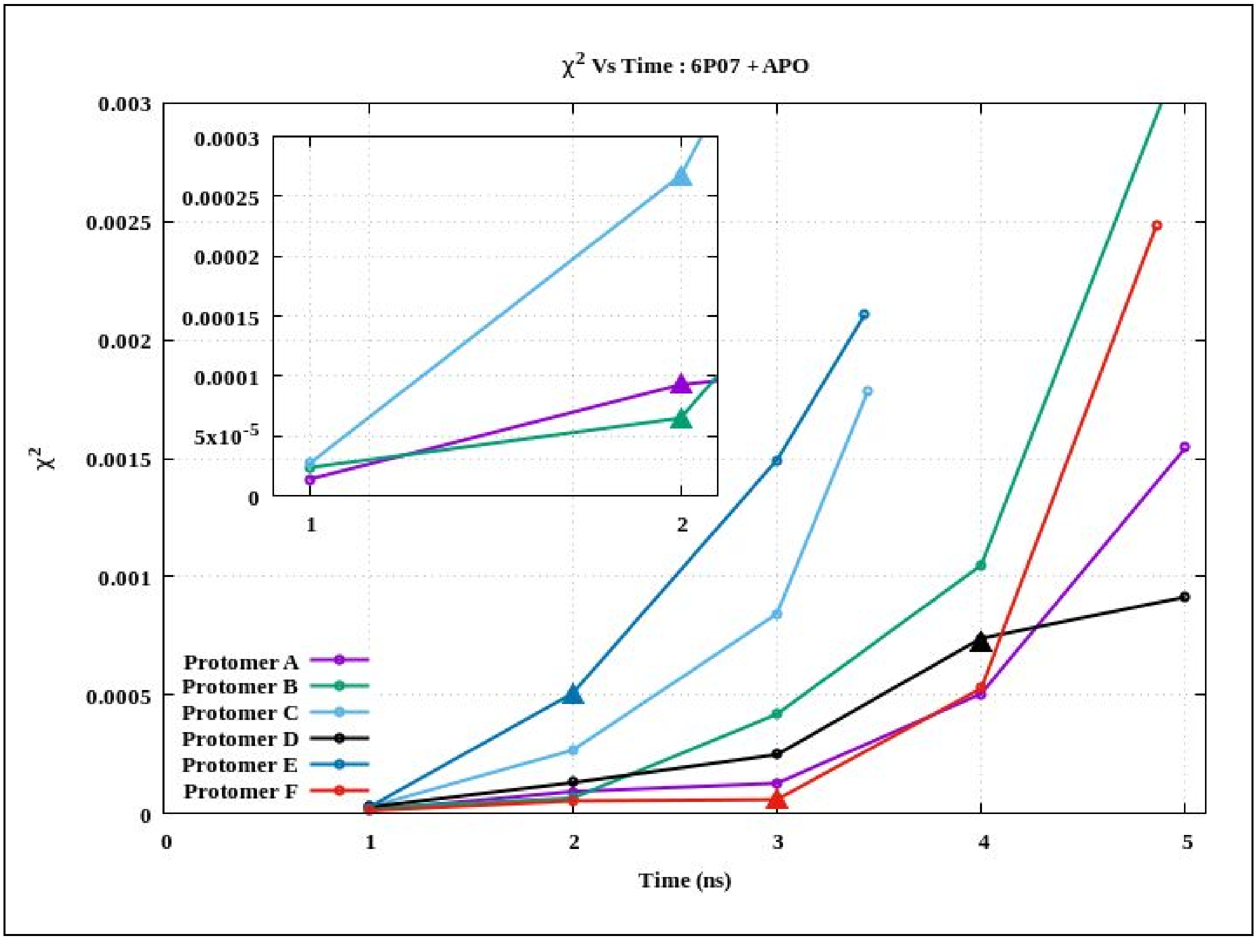


**Figure S14:** Representative **χ2** vs time (ns) plot showing elbow regions used for the selection of ***τ**** for each protomer in spastin, 6P07 APO state. The **χ2** values at elbow regions are denoted with "solid triangles".

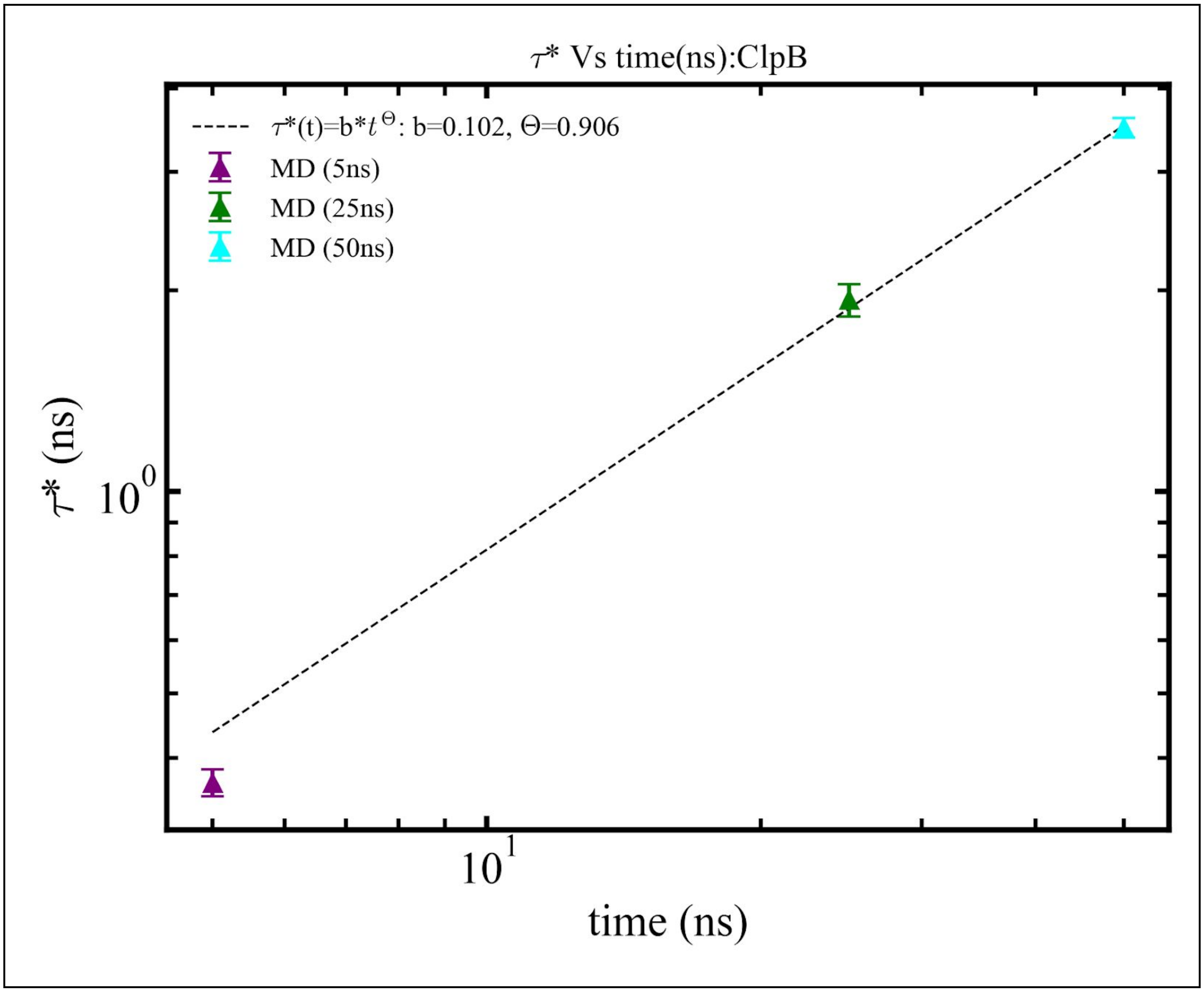


**Figure S15:** The $\tau$* vs time (ns) log-log plots showing the linear fitting for ClpB. The predicted $\tau$* values obtained at 1 millisecond timescale for ClpB is 28 μs. The dark magenta data points represent the values for the average of 6 protomers from the 5 ns trajectories, the dark green from the 25 ns trajectories, and the light cyan from the 50 ns trajectories and the error bars for the respective dots represent the standard error of $\tau$*.

#### References

[Sandate C, 2019] Sandate, Colby R., et al. "An allosteric network in spastin couples multiple activities required for microtubule severing." *Nat. Struct. Mol. Biol.* 26.8 (2019): 671-678.

[Eswar N, 2008] Eswar, Narayanan, et al., "Protein structure modeling with MODELLER." *Struct. prot.*. Humana Press, 2008. 145-159.

[Malde A, 2011] Malde, Alpeshkumar K., et al. "An automated force field topology builder (ATB) and repository: version 1.0." *J. Chem. Th. Comp.* 7.12 (2011): 4026-4037.

[Parrinello M, 1981] Parrinello, Michele, and Aneesur Rahman. "Polymorphic transitions in single crystals: A new molecular dynamics method." *J. Appl. Phys.* 52.12 (1981): 7182-7190.

[Daren T, 1993] Darden, Tom, Darrin York, and Lee Pedersen. "Particle mesh Ewald: An N · log (N) method for Ewald sums in large systems." *J. Chem. Phys.* 98.12 (1993): 10089-10092.

[Bussi G, 2007] Bussi, Giovanni, Davide Donadio, and Michele Parrinello. "Canonical sampling through velocity rescaling." *J. Chem. Phys.* 126.1 (2007): 014101.
